# Spatial distribution of Arctic bacterioplankton abundance is linked to distinct water masses and summertime phytoplankton bloom dynamics (Fram Strait, 79°N)

**DOI:** 10.1101/2020.09.02.277574

**Authors:** Magda G. Cardozo-Mino, Eduard Fadeev, Verena Salman-Carvalho, Antje Boetius

## Abstract

The Arctic is impacted by climate warming faster than any other oceanic region on Earth. Assessing the baseline of microbial communities in this rapidly changing ecosystem is vital for understanding the implications of ocean warming and sea ice retreat on ecosystem functioning. Using CARD-FISH and semi-automated counting, we quantified 14 ecologically relevant taxonomic groups of bacterioplankton (*Bacteria* and *Archaea*) from surface (0-30 m) down to deep waters (2500 m) in summerly ice-covered and ice-free regions of the Fram Strait, the main gateway for Atlantic inflow into the Arctic Ocean. Cell abundances of the bacterioplankton communities in surface waters varied from 10^5^ cells mL^-1^ in ice-covered regions to 10^6^ cells mL^-1^ in the ice-free regions, and were overall driven by variations in phytoplankton bloom conditions across the Strait. The bacterial classes *Bacteroidia* and *Gammaproteobacteria* showed several-fold higher cell abundances under late phytoplankton bloom conditions of the ice-free regions. Other taxonomic groups, such as the *Rhodobacteraceae,* revealed a distinct association of cell abundances with the surface Atlantic waters. With increasing depth (>500 m), the total cell abundances of the bacterioplankton communities decreased by up to two orders of magnitude, while largely unknown taxonomic groups (e.g., SAR324 and SAR202 clades) maintained constant cell abundances throughout the entire water column (ca. 10^3^ cells mL^-1^). This suggests that these enigmatic groups may occupy a specific ecological niche in the entire water column. Our results provide the first quantitative spatial variations assessment of bacterioplankton in the summerly ice-covered and ice-free Arctic water column, and suggest that further shift towards ice-free Arctic summers with longer phytoplankton blooms can lead to major changes in the associated standing stock of the bacterioplankton communities.

## 1 Introduction

Atmospheric and oceanic warming has a substantial impact on the Arctic Ocean already today (Dobricic et al., 2016; Sun et al., 2016; Dai et al., 2019). The strong decline in sea ice coverage (Peng and Meier, 2018; Dai et al., 2019) and heat transfer by the Atlantic water inflow (Beszczynska-Möller et al., 2012; Rudels et al., 2012; Walczowski et al., 2017) will affect stratification of the water column and can lead to an increase in upward mixing of the Atlantic core water, a process also termed “Atlantification” (Polyakov et al., 2017). The main inflow of Atlantic water into the Arctic Ocean occurs through the Fram Strait (Beszczynska-Möller et al., 2011), making it a sentinel region for observing the ongoing changes in the Arctic marine ecosystem (Soltwedel et al., 2005, 2016). The Fram Strait is also the main deep-water gateway between the Atlantic and the Arctic Ocean. It hosts two distinct hydrographic regimes; the West Spitsbergen Current (WSC) that carries relatively warm and saline Atlantic water northwards along the Svalbard shelf (Beszczynska-Möller et al., 2012; von Appen et al., 2015), and the East Greenland Current (EGC) that transports cold polar water and sea ice southwards from the Arctic Ocean along the ice-covered Greenland shelf (de Steur et al., 2009; Wekerle et al., 2017).

Sea ice conditions have a strong impact on the seasonal ecological dynamics in Fram Strait and the whole Arctic Ocean (Wassmann and Reigstad, 2011), affecting light availability and stratification in the water column. The presence of sea ice and snow cover can repress the seasonal phytoplankton bloom in the water column through light limitation (Mundy et al., 2005; Leu et al., 2011), or change its timing, e.g. by increasing stratification of the surface waters once the ice melts (Korhonen et al., 2013). Also, sea-ice algae can make up a significant proportion of the annual productivity (Leu et al., 2011; Boetius et al., 2013; Fernández-Méndez et al., 2014). Previous summer observations in the Fram Strait already suggested that total cell abundances and productivity of bacterioplankton communities in surface waters are driven by environmental parameters associated with phytoplankton bloom dynamics (Fadeev et al., 2018), such as the availability and composition of organic matter (Piontek et al., 2015; Engel et al., 2019), with differences between ice-covered and ice-free regions (Piontek et al., 2014; Fadeev et al., 2018).

Long-term summer observations in the region, conducted in the framework of the Long-Term Ecological Research (LTER) site HAUSGARTEN, revealed strong ecological variations associated with the Atlantic Meridional Overturning Circulation (AMOC; Soltwedel et al., 2016). Warming events during the past decades influenced seasonal phytoplankton blooms by causing a slow but continuous increase in biomass, and a shift from diatom- to flagellate-dominated communities (Nöthig et al., 2015; Engel et al., 2017; Basedow et al., 2018). It has been recently observed that phytoplankton blooms show an increasing partitioning of the produced organic carbon into the dissolved phase (Engel et al., 2019), which may result in a more active microbial loop in the upper ocean and less export of particulate matter (Vernet et al., 2017; Fadeev et al., 2020). In times of a rapidly changing Arctic ecosystem, investigating structure and dynamics of bacterioplankton communities remains a key component to the understanding of current changes in this environment. However, so far, an assessment of associated responses of the key bacterial taxa responsible for an increased recycling is missing, especially with regard to shifts in standing stocks.

To date, the majority of Arctic bacterioplankton studies are performed using high-throughput sequencing of the 16S rRNA gene, which cannot be directly converted to absolute standing stock abundances of specific taxonomic groups due to PCR and other quantitative biases (Gloor et al., 2017; Kumar et al., 2017; Piwosz et al., 2020). Here we used semi-automatic CAtalyzed Reporter Deposition-Fluorescence In Situ Hybridization (CARD-FISH; Pernthaler et al., 2002). The power of this technique lies in the ability to acquire absolute abundance of the targeted taxonomic groups free of compositional effect (Amann et al., 1990). Besides the ability to target and quantify specific taxonomic groups, the retrieval of a positive hybridization signal furthermore indicates that the analyzed cell was alive and active before fixation (Amann et al., 1990; DeLong et al., 1999). Automatization of the microscopic examination and counting procedure can reach a high-throughput standard (Schattenhofer et al., 2009; Teeling et al., 2012; Bižić-Ionescu et al., 2015; Bennke et al., 2016). Using CARD-FISH and semi-automated cell counting, we quantified cell abundances of 14 taxonomic groups in 44 samples, collected at 4 different water layers from surface (15-30 m) to the deep ocean (2500 m) in both, ice-free and ice-covered regions of the Fram Strait. The main objective of this study was to assess the standing stocks of key taxonomic groups in summerly Arctic bacterioplankton across the Strait. Using high-throughput data of bacterioplankton cell abundances we tested the following hypotheses: 1) in surface waters, the abundances of different bacterioplankton taxonomic groups are associated with phytoplankton bloom conditions, and are linked to the abundances of specific phytoplankton populations; 2) water depth structures the bacterioplankton communities, and 3) differences between communities in ice-covered and ice-free regions decrease with increasing water depth.

## 2 Results and Discussion

We investigated a total of 44 water samples from 11 stations that represented different hydrographic and biogeochemical conditions across the Fram Strait (Figure 1; Table S1). At each station, 4 different water layers were targeted: the surface mixed layer (0-30 m), epipelagic waters (100 m), deep mesopelagic waters (500-1000 m), and bathypelagic waters (1200-2500 m). Based on the known hydrography of the Strait (Rudels et al., 2012), and observed sea-ice conditions, the sampled stations were grouped into three distinct regions (Figure 1): the eastern ice-free region associated with the WSC (Beszczynska-Möller et al., 2012), the western ice-covered region associated with the EGC (de Steur et al., 2009), and the partially ice-covered region in the north-east that is associated with the highly productive ice-margin zone (further addressed as North- “N”) (Hebbeln and Wefer, 1991; Perrette et al., 2011). At the time of sampling in June-July 2016, microscopy counts of phytoplankton communities and chlorophyll *a* concentration, showed a late phytoplankton bloom across the Strait (Table S1; described in detail by Fadeev et al. 2020). The phytoplankton communities of the ice-covered EG and the ice-margin N stations were dominated by diatoms, in contrast to the ice-free HG stations that were dominated by *Phaeocystis* spp. These locally defined conditions correspond to an interannual trend of distinct phytoplankton bloom conditions observed in the western ice-covered EGC and the eastern ice-free WSC (Nöthig et al., 2015; Fadeev et al., 2018).

**Figure 1.**
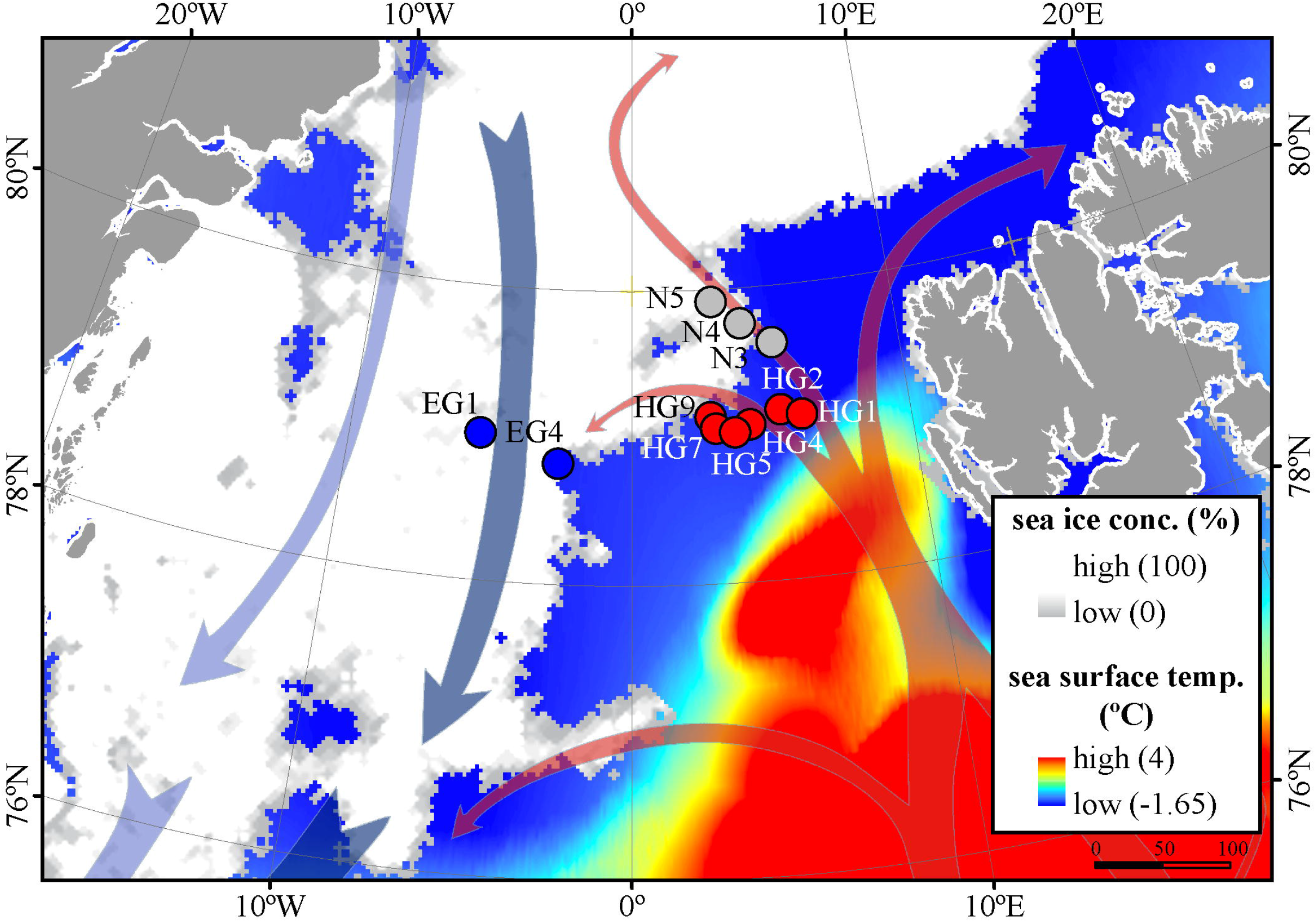
Oceanographic overview of the Fram Strait, including the monthly mean of sea-ice cover and sea surface temperature during July 2016. The sea ice concentration is represented by inverted grayscale (gray=low, white=high). Arrows represent general directions of the WSC (in red) and the EGC (in blue). Stations of water column sampling are indicated and colored according to their sea-ice conditions: ice-covered EGC stations - blue, ice-margin N stations - gray, ice-free WSC stations - red. The map was modified from (Fadeev et al., 2020).

### 2.1 Surface water bacterioplankton communities are affected by distinct phytoplankton bloom conditions

Phytoplankton blooms in surface waters generally lead to an increased cell abundance of heterotrophic bacteria that are specialized on degradation of organic matter from algal exudates and phytodetritus (Buchan et al., 2014; Teeling et al., 2016). Previous observations in the Fram Strait, acquired using high-throughput sequencing of the 16S rRNA gene, revealed a strong influence of the summerly phytoplankton bloom conditions on the bacterioplankton communities (Wilson et al., 2017; Müller et al., 2018b), differing between the ice-covered and ice-free regions of the Strait (Fadeev et al., 2018). During our sampling period, distinct phytoplankton bloom communities in surface waters across the Strait were observed, with a *Phaeocystis*-dominated bloom in the ice-free HG stations, a diatom-dominated bloom in the ice-covered EG stations, and mixed diatoms and *Phaeocystis* populations bloom in the ice-margin N stations (Fadeev et al., 2020). Along with this, we observed significantly higher cell abundances in the surface water total bacterioplankton communities of the HG and N stations (13-21×10^5^ cells mL^-1^), as compared to the EG stations (0.5-2×10^5^ cells mL^-1^; Kruskal-Wallis test; Chi square=81.85, df=2, *p*-value<0.01). The communities were dominated by bacterial cells that comprised 8-11×10^5^ cells mL^-1^ in the HG and N stations, and 2×10^5^ cells mL^-1^ in the EG stations. Within the bacterial communities, a combination of classes that are functionally associated with phytoplankton blooms in the region (*Bacteroidia*, *Gammaproteobacteria,* and the phylum *Verrucomicrobia*) (Fadeev et al., 2018) showed several-fold higher cell abundances in the HG and N stations (2.3-10×10^5^ cells mL^-1^), compared to the EG stations (0.5-1.5×10^5^ cells mL^-1^). Jointly, these classes comprised up to 50% of the analyzed bacterioplankton communities (Table 1). Other taxonomic groups, which were previously associated with the Arctic water masses and winter communities in the Fram Strait (e.g., the class *Deltaproteobacteria* and the *Thaumarchaeota*) (Wilson et al., 2017; Fadeev et al., 2018, 2020; Müller et al., 2018b), showed higher cell abundances in the ice-covered EG stations, as compared to the ice-free HG and ice-margin N stations (Table 1). Hence, the spatial variability in cell abundances of different taxonomic groups was apparently associated with different stages of the phytoplankton bloom and different water masses.

**Table 1.** Average cell abundances and proportions (% of DAPI stained cells) of selected taxonomic groups in surface (0-30 m) and epipelagic (100 m) depths of different regions across the Fram Strait. The proportions (%) were calculated based on the total bacterioplankton cell abundances, ‘n’ represents the number of counted fields of view. All values are represented in 10^5^ cells mL^-1^. SAR11 clade (SAR11), *Gammaproteobacteria* (GAM), *Alteromonadaceae/Colwelliaceae/Pseudoalteromonadaceae* (ATL), *Bacteroidetes* (BACT)*, Polaribacter* (POL)*, Verrucomicrobiales* (VER), *Opitutales* (OPI)*, Rhodobacteraceae* (ROS), *Deltaproteobacteria* (DELTA), and *Thaumarchaeota* (THA).

| Region | Water layer | SAR11 | % | n | GAM | % | n | ALT | % | n | BACT | % | n | POL | % | n |
| --- | --- | --- | --- | --- | --- | --- | --- | --- | --- | --- | --- | --- | --- | --- | --- | --- |
| EGC | Surface | 1.9 ± 0.7 | 50 | 53 | 0.3 ± 0.2 | 13 | 67 | 0.2 ± 0.1 | 7 | 64 | 0.6 ± 0.4 | 18 | 66 | 0.4 ± 0.3 | 11 | 62 |
| EGC | Epipelagic | 1.0 ± 0.6 | 26 | 50 | 0.2 ± 0.1 | 5 | 77 | 0.1 ± 0.0 | 2 | 61 | 0.3 ± 0.2 | 8 | 93 | 0.1 ± 0.0 | 2 | 52 |
| N | Surface | 4.2 ± 1.0 | 24 | 81 | 2.1 ± 0.6 | 13 | 78 | 0.1 ± 0.0 | 1 | 68 | 2.1 ± 0.8 | 12 | 85 | 1.3 ± 0.6 | 8 | 85 |
| N | Epipelagic | 2.2 ± 0.3 | 29 | 73 | 0.4 ± 0.1 | 5 | 109 | 0.2 ± 0.1 | 2 | 87 | 0.4 ± 0.1 | 6 | 102 | 0.1 ± 0.0 | 2 | 98 |
| WSC | Surface | 6.1 ± 2.0 | 38 | 150 | 1.6 ± 0.3 | 14 | 189 | 0.3 ± 0.1 | 2 | 186 | 2.1 ± 1.0 | 16 | 163 | 0.8 ± 0.2 | 7 | 162 |
| WSC | Epipelagic | 1.9 ± 0.2 | 32 | 175 | 0.2 ± 0.0 | 3 | 214 | 0.1 ± 0.0 | 1 | 186 | 0.2 ± 0.0 | 4 | 195 | 0.1 ± 0.0 | 1 | 176 |
| Region | Water layer | VER | % | n | OPI | % | n | ROS | % | n | DELTA | % | n | THA | % | n |
| EGC | Surface | 0.1 ± 0.0 | 3 | 41 | 0.1 ± 0.0 | 2 | 53 | 0.2 ± 0.1 | 7 | 44 | 0.2 ± 0.1 | 7 | 57 | 0.1 ± 0.0 | 3 | 34 |
| EGC | Epipelagic | 0.1 ± 0.0 | 2 | 63 | 0.1 ± 0.0 | 2 | 70 | 0.3 ± 0.2 | 9 | 60 | 0.1 ± 0.0 | 4 | 67 | 0.1 ± 0.02 | 4 | 75 |
| N | Surface | 0.9 ± 0.1 | 5 | 91 | 0.9 ± 0.01 | 5 | 91 | 0.6 ± 0.0 | 3 | 92 | 0.2 ± 0.1 | 1 | 86 | 0.1 ± 0.0 | 1 | 65 |
| N | Epipelagic | 0.2 ± 0.1 | 2 | 98 | 0.2 ± 0.1 | 2 | 107 | 0.2 ± 0.0 | 3 | 108 | 0.2 ± 0.0 | 3 | 94 | 0.3 ± 0.0 | 4 | 122 |
| WSC | Surface | 1.1 ± 0.0 | 5 | 171 | 0.8 ± 0.3 | 5 | 172 | 0.7 ± 0.3 | 4 | 165 | 0.1 ± 0.0 | 1 | 121 | 0.1 ± 0.0 | 1 | 178 |
| WSC | Epipelagic | 0.1 ± 0.0 | 2 | 219 | 0.1 ± 0.0 | 2 | 238 | 0.1 ± 0.0 | 2 | 223 | 0.2 ± 0.1 | 4 | 196 | 0.3 ± 0.1 | 5 | 246 |

To further test the link with distinct bloom conditions or distinct physical conditions in Atlantic vs. Arctic water masses, we conducted specific correlation tests between the cell abundances of different taxonomic groups, temperature and salinity (Table S2), which define the distinct water masses in the Fram Strait. We identified that cell abundances of *Verrucomicrobia* and its order *Opitutales,* as well as the SAR11 clade and the *Rhodobacteraceae* family (both members of the class *Alphaproteobacteria),* showed significant positive correlations to water temperature (Pearson’s correlation; *r*>0.5, *p*-value<0.05; Table S2), suggesting an association with the warmer Atlantic waters of the eastern Fram Strait. The *Verrucomicrobia* has been previously shown to be a major polysaccharide-degrading bacterial taxonomic group in Svalbard fjords (Cardman et al., 2014), and therefore may also be associated with the outflow from the Svalbard fjords into the Atlantic waters of the WSC (Cottier et al., 2005). The SAR11 clade and the *Rhodobacteraceae* have both been previously shown to correlate with temperature at high latitudes (Giebel et al., 2011; Tada et al., 2013), and are known to have distinct phylotypes in water masses with different temperatures (Selje et al., 2004; Sperling et al., 2012; Giovannoni, 2017). However, the *Rhodobacteraceae* are also known for their broad abilities in utilizing organic compounds (Buchan et al., 2014; Luo and Moran, 2014). Thus, one cannot rule out that their higher cell abundances in warmer waters of the HG and N stations are associated with the late stage of the phytoplankton bloom and their exudates. In addition, the SAR324 clade (*Deltaproteobacteria*) showed strong positive correlation with statistical significance to salinity (Pearson’s correlation; *r*>0.5, *p*-value<0.05; Table S2). During the summer, with increased melting of sea ice, a low-salinity water layer is formed in surface waters, and the strong stratification of this water layer enhances the development of the phytoplankton bloom (Fadeev et al., 2018). Consequently, the correlation of SAR324 with higher salinity suggests that their cell abundances are lower in surface waters where, in turn, we observe a strong phytoplankton bloom (e.g., in WSC).

The distinct surface water masses in the region differ not only in their physical but also in their biogeochemical characteristics (Wilson and Wallace, 1990; Fadeev et al., 2018), with higher concentrations of inorganic nitrogen and phosphate in the Atlantic, compared to the Arctic water masses. At the time of sampling, the typical Redfield ratio between inorganic nitrogen (mainly nitrate NO_3_) and inorganic phosphate (PO_4_) was below 16 (Redfield, 1963; Goldman et al., 1979). This suggests that the water masses were nitrogen limited across all three regions (Table S1) during summer due to phytoplankton dynamics. In order to disentangle the effect of biological consumption of nutrients from water mass-specific nutrient signatures, we calculated the seasonal net consumption of inorganic nutrients, as the proxy for phytoplankton bloom conditions (Table S1). Consumed nitrate (ΔNO_3_) and phosphate (ΔPO_4_) revealed a very strong positive correlation with statistical significance (Pearson’s correlation; *r*=0.86, *p*-value<0.05; Table S2). The consumed silica (ΔSiO_3_), used by diatoms, did not show a significant correlation to ΔPO_4_ and ΔNO_3_. This further supports the impact of different phytoplankton populations across the Strait (i.e., diatoms vs. *Phaeocystis;* Fadeev et al., 2020). Phytoplankton bloom-associated environmental parameters (chlorophyll *a* concentration and the consumed inorganic nutrients) revealed weaker relationships with cell abundances of different taxonomic groups (Table S2). Furthermore, we did not observe significant positive correlations of the cell abundances of diatoms or *Phaeocystis* spp., with the quantified bacterioplankton taxa. This might be explained by time lags and local differences in the dynamic development of phytoplankton blooms across the entire Strait (Wilson et al., 2017; Fadeev et al., 2018).

The complexity of Fram Strait surface waters with different ice-coverages, a dynamic ice-melt water layer and mesoscale mixing events of Atlantic and Polar water masses by eddies (Wekerle et al., 2017), challenges the identification of specific associations between microbial cell abundances and environmental parameters. While a mixture of all these environmental variables is likely shaping the bacterioplankton communities, our results showed that elevated cell abundances of some taxonomic groups (e.g., *Gammaproteobacteria*) had stronger association with phytoplankton bloom conditions observed at the site (e.g., through a link with algal exudates and nutrients as main source for growth) (Tada et al., 2011; Teeling et al., 2012). On the other hand, other taxonomic groups (e.g., SAR11 clade) were potentially more influenced by physical processes such as the presence of ice and distinct Arctic water masses (Kraemer et al., 2020).

### 2.2 Bacterioplankton communities strongly change in cell abundance and composition with depth

We found that in all three regions, total cell abundances of the entire bacterioplankton community were highest at surface with 10^5^-10^6^ cells mL^-1^, and significantly decreased with depth down to 10^4^ cells mL^-1^ at meso- and bathypelagic depths (Figure 2a; Table S3; Kruskal-Wallis test; Chi square=554.39, df=3,*p*-value<0.01). Members of the domain *Bacteria* dominated the communities throughout the entire water column, with highest cell abundances in surface waters (10^5^-10^6^ cells mL^-1^), and significantly lower 10^4^ cells mL^-1^ at depth (Figure 2b; Kruskal-Wallis test; Chi square=35.27, df=3, *p*-value<0.01). Archaeal cells had an overall lower abundance than bacterial cells by an order of magnitude throughout the entire water column, ranging from 10^4^ cells mL^-1^ at surface down to 10^3^ cells mL^-1^ in bathypelagic waters (Figure 2c). However, unlike *Bacteria*, archaeal communities doubled their absolute cell abundances from ca. 3×10^4^ cells mL^-1^ at surface to ca. 6×10^4^ cells mL^-1^ at 100 m depth, followed by a significant decrease in cell abundance at meso- and bathypelagic depths (Kruskal-Wallis test; Chi square=29.04, df=3, *p*-value<0.01). Compared to the stronger decline in bacterial cell numbers, this pattern mirrors the known global trend of relative archaeal enrichment in epipelagic waters (Karner et al., 2001; Herndl et al., 2005; Kirchman et al., 2007; Varela et al., 2008; Schattenhofer et al., 2009), and was also observed in other regions of the Arctic Ocean (Amano-Sato et al., 2013). Altogether, observed here bacterioplankton cell abundances in surface waters were well within the range of previous observations in the Fram Strait waters, conducted using flow cytometry (Piontek et al., 2014; Fadeev et al., 2018; Engel et al., 2019). However, compared to recent CARD-FISH based observations in eastern Fram Strait (Quero et al., 2020), cell abundances were consistently one order of magnitude lower along the entire water column. The discrepancy might be associated with methodological differences, such as shorter staining times and the usage of an automated over a manual counting approach in our study. Nevertheless, both studies showed a similar pattern of a strong decrease in bacterioplankton cell abundances with depth, which also matches observations in other oceanic regions (Karner et al., 2001; Church et al., 2003; Teira et al., 2004; Schattenhofer et al., 2009; Dobal-Amador et al., 2016).

**Figure 2.**
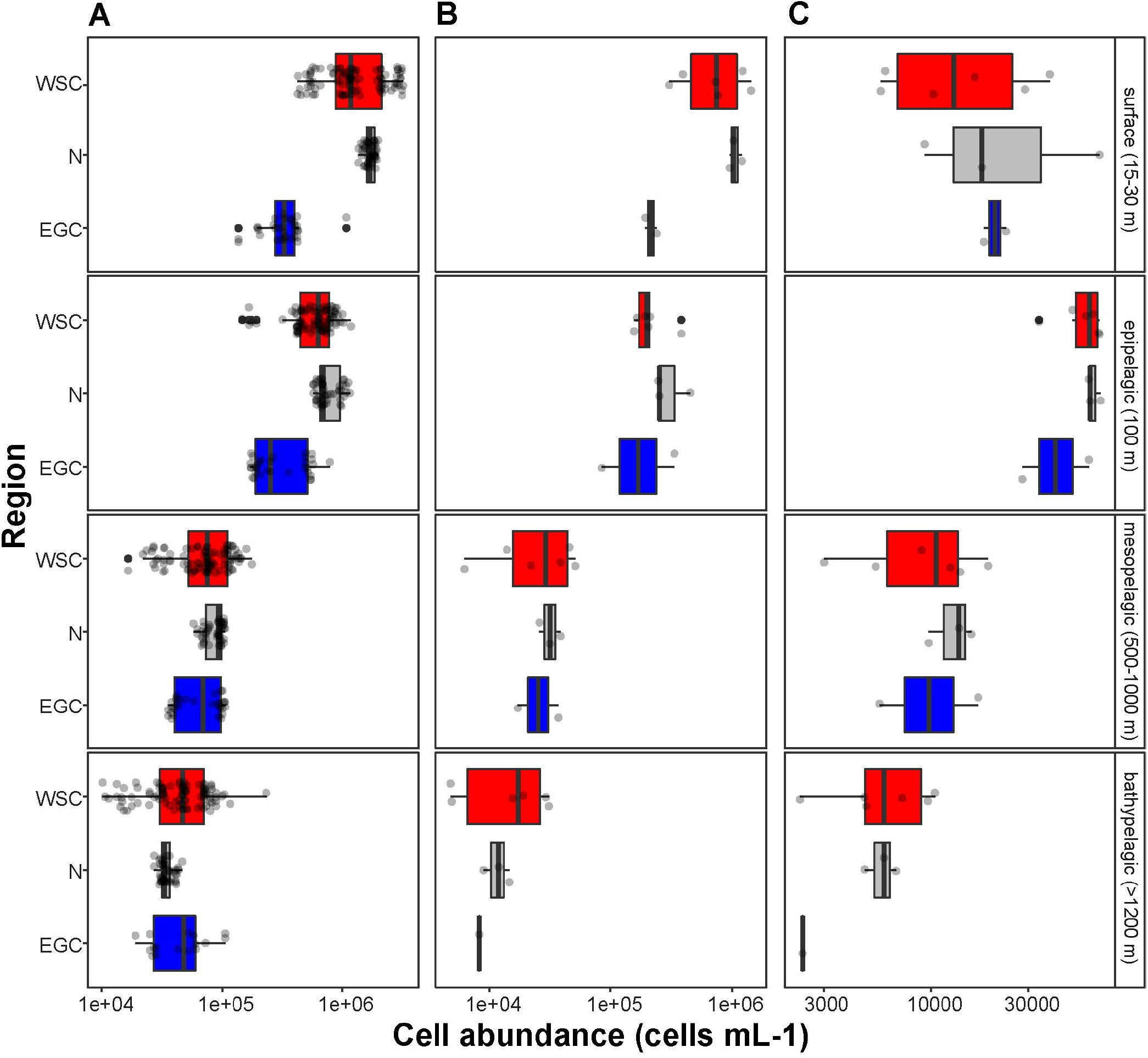
Bacterioplankton cell abundances in the different regions of the Fram Strait: Total bacterioplankton (A); *Bacteria* (B); and *Archaea* (C). Box plots were calculated based on cell abundance. Note the different scale of the cell abundances for *Archaea*. The different regions are indicated by color: ice-covered EGC-blue, ice-margin N - gray, ice-free WSC - red. The asterisks represent levels of statistical significance of difference between all three regions per depth and domain: * < 0.05, ** < 0.01, *** < 0.001.

In surface waters of all stations, ca. 60% of DAPI-stained total bacterioplankton community was covered by the *Bacteria*-specific probes (EUB388 I-III; Table S3). At depth (>100 m), the coverage of total cells by the *Bacteria*-specific probes strongly decreased to 16-40% of DAPI-stained cells (ANOVA; *F*_3_=15.39, *p*<0.01; Table S3). A similar decrease in detectability of the domain-specific probes was previously observed in other bacterioplankton microscopy studies (Karner et al., 2001; Herndl et al., 2005; Varela et al., 2008), and reasons may lie in a ribosomal nucleic acid concentration decrease within the bacterial cells (i.e., lower activity) towards the oligotrophic depths. In addition, there is a potential increase with greater water depths of microbial phylogenetic groups that are not captured by the currently existing probes (Hewson et al., 2006; Galand et al., 2009a; Agogué et al., 2011; Welch and Huse, 2011; Salazar et al., 2016).

Interestingly, the *Archaea*-specific probe (ARCH915) showed a different trend. In surface waters, the coverage of the probe was higher in the ice-covered EG stations (8% of DAPI-stained cells), compared to ice-free HG and ice-margin N stations (1-2% of DAPI-stained cells; Table S3). With depth (>100 m), in EG stations the coverage of the probe increased ca. twofold, while in HG and N stations the coverage of the probe increased ca. tenfold. Overall, across all three regions, coverage of the *Archaea*-specific probe was significantly higher at depth (ANOVA; *F*_3_=34.31, *p*<0.01), reaching 13-17% of DAPI-stained cells (Table S3). This trend implies an increase in relative abundance of *Archaea* with depth (Müller et al., 2018a; Fadeev et al., 2020). Taken together, our findings confirm previously observed higher abundances of *Archaea* in bacterioplankton communities of ice-covered waters (Wilson et al., 2017; Müller et al., 2018b; Fadeev et al., 2020), and correspond to the globally observed trend of an increasing archaeal importance at depth (Herndl et al., 2005; Teira et al., 2006; Galand et al., 2009b).

### 2.3 Enigmatic microbial lineages increase in cell abundance towards the deep ocean

The deep waters of the Fram Strait basin (>500 m) have a rather homogeneous hydrography (von Appen et al., 2015), and are less affected by the seasonal dynamics that govern the surface layers (Wilson et al., 2017). Previous molecular observations of the deep water bacterioplankton communities showed high sequence abundances of largely unknown taxonomic groups, such as the SAR202 (class *Dehalococcoidia),* SAR324 (*Deltaproteobacteria),* and SAR406 (phylum *Marinimicrobia*) (Wilson et al., 2017; Fadeev et al., 2020; Quero et al., 2020). There was also higher archaeal sequence abundance at depth, with the class *Nitrososphaeria* reaching up to 15% of the sequences in mesopelagic waters (> 200 m) (Wilson et al., 2017; Müller et al., 2018b; Fadeev et al., 2020). However, it has also been recently shown that in ice-covered regions of the Strait surface-dominant taxonomic groups, such as *Gammaproteobacteria* and *Nitrososphaeria*, are exported via fast-sinking aggregates from surface to the deep ocean (>1000 m), where they may realize an ecological niche (Fadeev et al., 2020). We observed that in all meso- and bathypelagic waters across all analyzed regions the total cell abundances of the bacterioplankton communities were in the range of 10^4^ cells mL^-1^ (Figure 2), reflecting observations made in other regions of the Arctic Ocean (Wells and Deming, 2003; Wells et al., 2006). Bacterial taxonomic groups that dominated the surface water communities (e.g., *Bacteroidetes, Gammaproteobacteria* and *Verrucomicrobia),* in both ice-free and ice-covered regions of the Strait, decreased by two orders of magnitude in their cell abundances at meso- and bathypelagic depths (Kruskal-Wallis test; *p*-value<0.01; Table 2). This trend strongly correlated with the total bacterioplankton cell abundances along the general water column (Pearson’s correlation; *r*>0.8, *p*-value<0.05; Figure S1). In contrast, other bacterial groups, such as the SAR202 and SAR324 clades, proportionally increased in cell abundances with depth, and maintained overall constant cell abundances of ca. 0.5×10^4^ cells mL^-1^ until the deep basin (Table 2). Previous molecular studies of bacterioplankton communities in the Fram Strait suggested a proportional increase of these largely understudied bacterial lineages in the deep ocean, which were previously found to be associated with winterly (surface) bacterioplankton (Wilson et al., 2017; Fadeev et al., 2020). The cell abundances presented here indicate that their increasing proportional abundance at depth is due to stronger decrease in the cell abundances of other groups (Table 2; Table S4). Very little is currently known about these two taxonomic groups, but previous genetic observations suggest that they possess distinct metabolic capabilities, and may be involved in the degradation of recalcitrant organic matter (SAR202 clade; Landry et al., 2017; Colatriano et al., 2018; Saw et al., 2019), or, in sulfur oxidation (SAR324 clade; Swan et al., 2011; Sheik et al., 2014). Their homogeneous distribution from the stratified surface to the homogenous deep ocean of the Fram Strait suggests that these enigmatic bacterial groups fulfil an ecological niche that exists in the entire water column, and thus may have unique roles in oceanic nutrient cycling.

**Table 2.** Average cell abundances and proportions (% of DAPI stained cells) of selected taxonomic groups in deep water layers of the different regions across the Fram Strait. The proportions (%) were calculated based on the total bacterioplankton cell abundances, ‘n’ represents the number of counted fields of view. Standard error was not calculated for samples of the EGC located in the bathypelagic zone due to one station located at this depth in the region. All values are represented in 10^5^ cells mL^-1^. *Chloroflexi* (CFX)*, Deltaproteobacteria* (DELTA)*, Thaumarchaeota* (THA), *Rhodobacteraceae* (ROS), SAR202, SAR324, SAR406 and SAR11 clades.

| Region | Water layer | CFX | % | n | DELTA | % | n | THA | % | n | ROS | % | n |
| --- | --- | --- | --- | --- | --- | --- | --- | --- | --- | --- | --- | --- | --- |
| EGC | Mesopelagic | 0.02 ± 0.0 | 5 | 57 | 0.03 ± 0.01 | 5 | 45 | 0.01 ± 0.0 | 2 | 27 | 0.1 ± 0.0 | 11 | 75 |
| EGC | Bathypelagic | 0.02 ± NA | 4 | 18 | 0.02 ± NA | 3 | 33 | 0.01 ± NA | 2 | 14 | 0.03 ± NA | 5 | 14 |
| N | Mesopelagic | 0.03 ± 0.0 | 3 | 116 | 0.04 ± 0.0 | 5 | 99 | 0.01 ± 0.0 | 1 | 28 | 0.1 ± 0.01 | 13 | 128 |
| N | Bathypelagic | 0.02 ± 0.0 | 6 | 115 | 0.03 ± 0.0 | 8 | 123 | 0.01 ± 0.0 | 4 | 39 | 0.1 ± 0.0 | 20 | 122 |
| WSC | Mesopelagic | 0.03 ± 0.0 | 3 | 217 | 0.1 ± 0.01 | 6 | 139 | 0.02 ± 0.0 | 3 | 86 | 0.1 ± 0.01 | 7 | 195 |
| WSC | Bathypelagic | 0.02 ± 0.0 | 4 | 199 | 0.1 ± 0.02 | 9 | 130 | 0.02 ± 0.0 | 4 | 54 | 0.04 ± 0.01 | 14 | 197 |
| Region | Water layer | SAR202 | % | n | SAR324 | % | n | SAR406 | % | n | SAR11 | % | n |
| EGC | Mesopelagic | 0.02 ± 0.0 | 4 | 55 | 0.05 ± 0.0 | 6 | 60 | 0.01 ± 0.0 | 1 | 11 | 0.15 ± 0.1 | 22 | 49 |
| EGC | Bathypelagic | 0.02 ± NA | 5 | 26 | 0.03 ± NA | 5 | 35 | 0.01 ± NA | 3 | 6 | 0.07 ± NA | 14 | 32 |
| N | Mesopelagic | 0.03 ± 0.0 | 3 | 107 | 0.06 ± 0.0 | 6 | 66 | 0.01 ± 0.0 | 1 | 18 | 0.17 ± 0.0 | 20 | 101 |
| N | Bathypelagic | 0.03 ± 0.0 | 8 | 120 | 0.03 ± 0.0 | 8 | 54 | 0.01 ± 0.0 | 3 | 37 | 0.09 ± 0.0 | 27 | 85 |
| WSC | Mesopelagic | $0.03 \pm 0.0$ | 5 | 215 | $0.05 \pm 0.0$ | 5 | 168 | $0.01 \pm 0.0$ | 2 | 37 | $0.16 \pm 0.0$ | 19 | 166 |
| WSC | Bathypelagic | $0.03 \pm 0.0$ | 7 | 210 | $0.05 \pm 0.0$ | 6 | 138 | $0.01 \pm 0.0$ | 3 | 45 | $0.09 \pm 0.0$ | 21 | 201 |

The proportional decrease of archaeal cell abundances with depth was less than that of members of the domain *Bacteria* (Table S4), meaning that members of the *Archaea* were proportionally increasing in the total microbial deep-water communities. The *Thaumarchaeota* strongly correlated with the pattern of the archaeal cell abundances (Pearson’s correlation; *r*=0.76, *p*-value<0.05; Figure S1), showing a two-fold increase in cell abundance from surface to epipelagic depth (100 m), followed by a substantial decrease towards meso- and bathypelagic waters (Table S4). This two-fold increase towards the epipelagic depths corresponds to previous observations of *Thaumarchaeota* in the north Atlantic (Müller et al., 2018b) and further increase in cell abundances at higher depths (>1000 m) was also observed in other oceanic regions (Karner et al., 2001; Church et al., 2003; Herndl et al., 2005). It has been shown in molecular studies that *Thaumarchaeota* comprise a large proportion of the bacterioplankton communities in the Fram Strait, especially in the epipelagic waters (Wilson et al., 2017; Müller et al., 2018b; Fadeev et al., 2020). In our study, the *Thaumarchaeota* exhibited their highest cell abundances at 100 m in the ice-free HG, and at the ice-margin N stations (3×10^4^ cells mL^-1^), where they comprised half of the total archaeal community (Table S4). The strong absolute decrease of *Thaumarchaeota* cell abundances towards the meso- and bathypelagic waters suggests a decrease in activity with depth (Herndl et al., 2005; Kirchman et al., 2007; Alonso-Sáez et al., 2012), and thus lower cell detectability. In deeper water layers, other pelagic archaeal groups, such as the phylum *Euryarchaeota* that was not quantified in this study, may increase in abundance and form the bulk of total archaeal cells here (Galand et al., 2010; Fadeev et al., 2020).

## 3 Conclusions

Using state-of-the-art semi-automatic microscopy cell counting, we quantified the absolute cell abundance of 14 key taxonomic groups in summer bacterioplankton communities of the Fram Strait. Our observations covered both the ice-free and ice-covered regions of the Strait, which at the time of sampling were characterized by different phytoplankton bloom stages. Our results showed that in surface waters, abundance of some taxonomic groups was related to the Atlantic waters (e.g., *Rhodobacteraceae*). The abundance of different taxonomic groups was strongly positively (e.g., *Gammaproteobacteria*) and negatively (e.g., SAR324 clade) associated with the states of the seasonal phytoplankton bloom across the Strait. Based on previous studies in the region, it is conceivable that there were also specific associations between the of blooming phytoplankton and different bacterioplankton taxa, however these were not observed in our analysis. This suggests that currently predicted longer seasonal phytoplankton blooms, as well as the increasing Atlantic influence on the Arctic Ocean (i.e., ‘Atlantification’), may have a strong impact on the composition and biogeographical distribution of certain bacterioplankton taxonomic groups in the surface Arctic waters.

This study also provides the first extensive quantification of bacterioplankton communities in the deep Arctic water column (> 500 m). We showed that with depth, some taxonomic groups, such as the SAR202 clade, maintained similar abundances throughout the entire water column (2500 m depth), where other taxa decline by several-fold. This observation suggests that despite their low abundance, some taxonomic groups may potentially realize a unique ecological niche throughout the entire water column.

Altogether, our quantitative data on cell abundances of ecologically relevant taxonomic bacterioplankton groups provide insight into factors structuring pelagic bacterioplankton communities from surface to the deep waters of the Arctic Ocean and a baseline to better assess future changes in a rapidly warming region.

## 4 Materials and Methods

### 4.1 Sampling and environmental data collection

Sampling was carried out during the RV Polarstern expedition PS99.2 to the Long-Term Ecological Research (LTER) site HAUSGARTEN in Fram Strait (June 24th – July 16th, 2016). Sampling was carried out with 12 L Niskin bottles mounted on a CTD rosette (Sea-Bird Electronics Inc. SBE 911 plus probe) equipped with temperature and conductivity sensors, a pressure sensor, altimeter, and a chlorophyll fluorometer. On board, the samples were fixed with formalin in a final concentration of 2% for 10 – 12 hours, then filtered onto 0.2 μm polycarbonate Nucleopore Track-Etched filters (Whatman, Buckinghamshire, UK), and stored at −20°C for further analysis.

Hydrographic data of the seawater including temperature and salinity were retrieved from PANGAEA (Schröder and Wisotzki, 2014), along with measured chlorophyll *a* concentration (Nöthig et al., 2018; Fadeev et al., 2020) (Table S1).

### 4.2 Catalyzed reporter deposition-fluorescence in situ hybridization (CARD-FISH)

We quantified absolute cell abundances of 14 key bacterioplankton groups (Table S5), based on their relatively high sequence abundance and recurrences in previous molecular studies of Arctic waters (Bowman et al., 2012; Wilson et al., 2017; Müller et al., 2018b; Fadeev et al., 2020). CARD-FISH was applied based on the protocol established by (Pernthaler et al., 2002), using horseradish-peroxidase (HRP)–labelled oligonucleotide probes (Biomers.net, Ulm, Germany). All probes were checked for specificity and coverage of their target groups against the SILVA database release 132 (Quast et al., 2013). All filters were embedded in 0.2% low-gelling-point agarose, and treated with 10 mg mL^-1^ lysozyme solution (Sigma-Aldrich Chemie GmbH, Hamburg, Germany) for 1 h at 37°C. Filters for enumerating *Archaea* and *Thaumarchaeota* were treated for an additional 30 min in 36 U mL^-1^ achromopeptidase (Sigma-Aldrich Chemie GmbH, Hamburg, Germany) and 15 μg mL^-1^ proteinase K at 37°C. Subsequently, endogenous peroxidases were inactivated by submerging the filter pieces in 0.15% H_2_O_2_ in methanol for 30 min before rinsing in Milli-Q water and dehydration in 96% ethanol. Then, the filters were covered in hybridization buffer and a probe concentration of 0.2 ng μL^-1^. Hybridization was performed at 46°C for 2.5 h, followed by washing in pre-warmed washing buffer at 48°C for 10 min, and 15 min in 1x PBS. Signal amplification was carried out for 45 min at 46°C with amplification buffer containing either tyramide-bound Alexa 488 (1 μg/mL) or Alexa 594 (0.33 μg mL^-1^). Afterwards, the cells were counterstained in 1 μg/mL DAPI (4’,6-diamidino-2-phenylindole; Thermo Fisher Scientific GmbH, Bremen, Germany) for 10 min at 46°C. After rinsing with Milli-Q water and 96% ethanol, the filter pieces were embedded in a 4:1 mix of Citifluor (Citifluor Ltd, London, United Kingdom) and Vectashield (Vector Laboratories, Inc., Burlingame, United States), and stored overnight at −20°C for later microscopy evaluation.

### 4.3 Automated image acquisition and cell counting

The filters were evaluated microscopically under a Zeiss Axio Imager.Z2 stand (Carl Zeiss MicroImaging GmbH, Jena, Germany), equipped with a multipurpose fully automated microscope imaging system (MPISYS), a Colibri LED light source illumination system, and a multi-filter set 62HE (Carl Zeiss MicroImaging GmbH, Jena, Germany). Pictures were taken via a cooled charged-coupled-device (CCD) camera (AxioCam MRm; Carl Zeiss AG, Oberkochen, Germany) with a 63× oil objective, a numerical aperture of 1.4, and a pixel size of 0.1016 μm/pixel, coupled to the AxioVision SE64 Rel.4.9.1 software (Carl Zeiss AG, Oberkochen, Germany) as described by (Bennke et al., 2016). Exposure times were adjusted after manual inspection with the AxioVision Rel.4.8 software coupled to the SamLoc 1.7 software (Zeder et al., 2011), which was also used to define the coordinates of the filters on the slides. For image acquisition, channels were defined with the MPISYS software, and a minimum of 55 fields of view with a minimum distance of 0.25 mm were acquired of each filter piece by recoding a z-stack of 7 images in autofocus.

Cell enumeration was performed with the software Automated Cell Measuring and Enumeration Tool (ACMETool3, 2018-11-09; M. Zeder, Technobiology GmbH, Buchrain, Switzerland). Cells were counted as objects according to manually defined parameters separately for the DAPI and FISH channels.

### 4.4 Calculation of consumed inorganic nutrients

Following (Fadeev et al., 2018) the nutrient consumption (Δ) at each station was calculated by subtracting the mean value of all collected measurements above 50 m from the mean value of all collected measurements between 50 and 100 m (below the seasonal pycnocline).

### 4.5 Statistical analyses

All statistical analyses and calculations in this study were performed using R (v4.0.2) (www.r-project.org) in RStudio (v1.3.1056), *i.e.* statistical tests for normality, ANOVA and Kruskal-Wallis. Post-hoc Wilcoxon test and Pearson’s rank correlation coefficient were conducted with the R package “rstatix” (v0.6.0) (Kassambara, 2020). Plots were generated using the R package “ggplot2” (v3.3.2) (Wickham, 2016) and “tidyverse” (v1.3.0) (Wickham et al., 2019).

### 4.6 Data availability

All data is accessible via the Data Publisher for Earth & Environmental Science PANGAEA (www.pangaea.de): cell abundances under doi:10.1594/PANGAEA.905212, and inorganic nutrient measurements under doi:10.1594/PANGAEA.906132. Scripts for processing the data can be accessed at https://github.com/edfadeev/FramStrait-counts.

## 5 Author contributions

MC-M, EF and VS-C designed and conducted the study. MC-M, EF and VS-C wrote the manuscript with guidance from AB. All authors critically revised the manuscript and gave their approval of the submitted version.

## 7 Acknowledgements

We thank the captain and crew of RV Polarstern expedition PS99.2, as well as the chief scientist Thomas Soltwedel for support with work at sea. We also thank Pier Offre for assistance in sampling, Greta Reintjes for designing the probe Opi346, Mareike Bach for technical support, Sinhue Torres-Valdes and Laura Wischnewski for conducting the inorganic nutrient measurements. We thank Andreas Ellrott for the support with the automated microscope from the Max Planck Institute for Marine Microbiology. This work was conducted in the framework of the HGF Infrastructure Program FRAM of the Alfred-Wegener-Institute Helmholtz Center for Polar and Marine.

## References

Agogué, H., Lamy, D., Neal, P. R., Sogin, M. L., and Herndl, G. J. (2011). Water mass-specificity of bacterial communities in the North Atlantic revealed by massively parallel sequencing. Mol. Ecol. 20, 258–274. doi:10.1111/j.1365-294X.2010.04932.x.

Alonso-Sáez, L., Waller, A. S., Mende, D. R., Bakker, K., Farnelid, H., Yager, P. L., et al. (2012). Role for urea in nitrification by polar marine Archaea. Proc. Natl. Acad. Sci. U. S. A. 109, 17989–17994. doi:10.1073/pnas.1201914109.

Amann, R. I., Binder, B. J., Olson, R. J., Chisholm, S. W., Devereux, R., and Stahl, D. A. (1990). Combination of 16S rRNA-targeted oligonucleotide probes with flow cytometry for analyzing mixed microbial populations. Appl. Environ. Microbiol. 56, 1919–1925. doi:10.1128/aem.56.6.1919-1925.1990.

Amano-Sato, C., Akiyama, S., Uchida, M., Shimada, K., and Utsumi, M. (2013). Archaeal distribution and abundance in water masses of the Arctic Ocean, Pacific sector. Aquat. Microb. Ecol. 69, 101–112. doi:10.3354/ame01624.

Basedow, S. L., Sundfjord, A., von Appen, W. J., Halvorsen, E., Kwasniewski, S., and Reigstad, M. (2018). Seasonal variation in transport of Zooplankton Into the Arctic basin through the Atlantic Gateway, Fram Strait. Front. Mar. Sci. 5, 194. doi:10.3389/fmars.2018.00194.

Bennke, C. M., Reintjes, G., Schattenhofer, M., Ellrott, A., Wulf, J., Zeder, M., et al. (2016). Modification of a high-throughput automatic microbial cell enumeration system for shipboard analyses. Appl. Environ. Microbiol. 82, 3289–3296. doi:10.1128/AEM.03931-15.

Beszczynska-Möller, A., Fahrbach, E., Schauer, U., and Hansen, E. (2012). Variability in Atlantic water temperature and transport at the entrance to the Arctic Ocean, 19972010. ICES J. Mar. Sci. 69, 852–863. doi:10.1093/icesjms/fss056.

Beszczynska-Möller, A., Woodgate, R. A., Lee, C., Melling, H., and Karcher, M. (2011). A synthesis of exchanges through the main oceanic gateways to the Arctic Ocean. Oceanography 24, 83–99. doi:10.5670/oceanog.2011.59.

Bižić-Ionescu, M., Zeder, M., Ionescu, D., Orlić, S., Fuchs, B. M., Grossart, H. P., et al. (2015). Comparison of bacterial communities on limnic versus coastal marine particles reveals profound differences in colonization. Environ. Microbiol. 17, 3500–3514. doi:10.1111/1462-2920.12466.

Boetius, A., Albrecht, S., Bakker, K., Bienhold, C., Felden, J., Fernández-Méndez, M., et al. (2013). Export of algal biomass from the melting arctic sea ice. Science (80-.). 339, 1430–1432. doi:10.1126/science.1231346.

Bowman, J. S., Rasmussen, S., Blom, N., Deming, J. W., Rysgaard, S., and Sicheritz-Ponten, T. (2012). Microbial community structure of Arctic multiyear sea ice and surface seawater by 454 sequencing of the 16S RNA gene. ISME J. 6, 11–20. doi:10.1038/ismej.2011.76.

Buchan, A., LeCleir, G. R., Gulvik, C. A., and González, J. M. (2014). Master recyclers: features and functions of bacteria associated with phytoplankton blooms. Nat. Rev. Microbiol. 12, 686–698. doi:10.1038/nrmicro3326.

Cardman, Z., Arnosti, C., Durbin, A., Ziervogel, K., Cox, C., Steen, A. D., et al. (2014). Verrucomicrobia are candidates for polysaccharide-degrading bacterioplankton in an Arctic fjord of Svalbard. Appl. Environ. Microbiol. 80, 3749–3756. doi:10.1128/AEM.00899-14.

Church, M. J., DeLong, E. F., Ducklow, H. W., Karner, M. B., Preston, C. M., and Karl, D. M. (2003). Abundance and distribution of planktonic Archaea and Bacteria in the waters west of the Antarctic Peninsula. Limnol. Oceanogr. 48, 1893–1902. doi:10.4319/lo.2003.48.5.1893.

Colatriano, D., Tran, P. Q., Guéguen, C., Williams, W. J., Lovejoy, C., and Walsh, D. A. (2018). Genomic evidence for the degradation of terrestrial organic matter by pelagic Arctic Ocean Chloroflexi bacteria. Commun. Biol. 1, 90. doi:10.1038/s42003-018-0086-7.

Cottier, F., Tverberg, V., Inall, M., Svendsen, H., Nilsen, F., and Griffiths, C. (2005). Water mass modification in an Arctic fjord through cross-shelf exchange: The seasonal hydrography of Kongsfjorden, Svalbard. J. Geophys. Res. Ocean. 110, 1–18. doi:10.1029/2004JC002757.

Dai, A., Luo, D., Song, M., and Liu, J. (2019). Arctic amplification is caused by sea-ice loss under increasing CO_2_. Nat. Commun. 10, 121. doi:10.1038/s41467-018-07954-9.

de Steur, L., Hansen, E., Gerdes, R., Karcher, M., Fahrbach, E., and Holfort, J. (2009). Freshwater fluxes in the East Greenland Current: A decade of observations. Geophys. Res. Lett. 36, L23611. doi:10.1029/2009GL041278.

DeLong, E. F., Taylor, L. T., Marsh, T. L., and Preston, C. M. (1999). Visualization and enumeration of marine planktonic archaea and bacteria by using polyribonucleotide probes and fluorescent in situ hybridization. Appl. Environ. Microbiol. 65, 5554–5563. doi:10.1128/aem.65.12.5554-5563.1999.

Dobal-Amador, V., Nieto-Cid, M., Guerrero-Feijoo, E., Hernando-Morales, V., Teira, E., and Varela-Rozados, M. M. (2016). Vertical stratification of bacterial communities driven by multiple environmental factors in the waters (0-5000 m) off the Galician coast (NW Iberian margin). Deep. Res. Part I Oceanogr. Res. Pap. 114, 1–11. doi:10.1016/j.dsr.2016.04.009.

Dobricic, S., Vignati, E., and Russo, S. (2016). Large-scale atmospheric warming in winter and the arctic sea ice retreat. J. Clim. 29, 2869–2888. doi:10.1175/JCLI-D-15-0417.1.

Engel, A., Bracher, A., Dinter, T., Endres, S., Grosse, J., Metfies, K., et al. (2019). Inter-annual variability of organic carbon concentrations in the eastern Fram Strait during summer (2009-2017). Front. Mar. Sci. 6, 187. doi:10.3389/fmars.2019.00187.

Engel, A., Piontek, J., Metfies, K., Endres, S., Sprong, P., Peeken, I., et al. (2017). Inter-annual variability of transparent exopolymer particles in the Arctic Ocean reveals high sensitivity to ecosystem changes. Sci. Rep. 7, 4129. doi:10.1038/s41598-017-04106-9.

Fadeev, E., Rogge, A., Ramondenc, S., Nöthig, E.-M., Wekerle, C., Bienhold, C., et al. (2020). Sea-ice retreat may decrease carbon export and vertical microbial connectivity in the Eurasian Arctic basins. Nat. Res. doi:10.21203/rs.3.rs-101878/v1.

Fadeev, E., Salter, I., Schourup-Kristensen, V., Nöthig, E. M., Metfies, K., Engel, A., et al. (2018). Microbial communities in the east and west fram strait during sea ice melting season. Front. Mar. Sci. 5, 429. doi:10.3389/fmars.2018.00429.

Fernández-Méndez, M., Wenzhöfer, F., Peeken, I., Sørensen, H. L., Glud, R. N., and Boetius, A. (2014). Composition, buoyancy regulation and fate of ice algal aggregates in the Central Arctic Ocean. PLoS One 9, e107452–e107452. doi:10.1371/journal.pone.0107452.

Galand, P. E., Casamayor, E. O., Kirchman, D. L., and Lovejoy, C. (2009a). Ecology of the rare microbial biosphere of the Arctic Ocean. Proc. Natl. Acad. Sci. U. S. A. 106, 22427–22432. doi:10.1073/pnas.0908284106.

Galand, P. E., Casamayor, E. O., Kirchman, D. L., Potvin, M., and Lovejoy, C. (2009b). Unique archaeal assemblages in the arctic ocean unveiled by massively parallel tag sequencing. ISME J. 3, 860–869. doi:10.1038/ismej.2009.23.

Galand, P. E., Potvin, M., Casamayor, E. O., and Lovejoy, C. (2010). Hydrography shapes bacterial biogeography of the deep Arctic Ocean. ISME J. 4, 564–576. doi:10.1038/ismej.2009.134.

Giebel, H. A., Kalhoefer, D., Lemke, A., Thole, S., Gahl-Janssen, R., Simon, M., et al. (2011). Distribution of Roseobacter RCA and SAR11 lineages in the North Sea and characteristics of an abundant RCA isolate. ISME J. 5, 8–19. doi:10.1038/ismej.2010.87.

Giovannoni, S. J. (2017). SAR11 Bacteria: The Most Abundant Plankton in the Oceans. Ann. Rev. Mar. Sci. 9, 231–255. doi:10.1146/annurev-marine-010814-015934.

Gloor, G. B., Macklaim, J. M., Pawlowsky-Glahn, V., and Egozcue, J. J. (2017). Microbiome datasets are compositional: And this is not optional. Front. Microbiol. 8, 2224. doi:10.3389/fmicb.2017.02224.

Goldman, J. C., McCarthy, J. J., and Peavey, D. G. (1979). Growth rate influence on the chemical composition of phytoplankton in oceanic waters. Nature 279, 210–215. doi:10.1038/279210a0.

Hebbeln, D., and Wefer, G. (1991). Effects of ice coverage and ice-rafted material on sedimentation in the Fram Strait. Nature 350, 409–411. doi:10.1038/350409a0.

Herndl, G. J., Reinthaler, T., Teira, E., Van Aken, H., Veth, C., Pernthaler, A., et al. (2005). Contribution of Archaea to total prokaryotic production in the deep atlantic ocean. Appl. Environ. Microbiol. 71, 2303–2309. doi:10.1128/AEM.71.5.2303-2309.2005.

Hewson, I., Steele, J. A., Capone, D. G., and Fuhrman, J. A. (2006). Remarkable heterogeneity in meso- and bathypelagic bacterioplankton assemblage composition. Limnol. Oceanogr. 51, 1274–1283. doi:10.4319/lo.2006.51.3.1274.

Karner, M. B., Delong, E. F., and Karl, D. M. (2001). Archaeal dominance in the mesopelagic zone of the Pacific Ocean. Nature 409, 507–510. doi:10.1038/35054051.

Kassambara, A. (2020). rstatix: Pipe-friendly framework for basic statistical tests. R package version 0.5.0.999. R Packag. version 0.6.0, https://rpkgs.datanovia.com/rstatix/.

Kirchman, D. L., Elifantz, H., Dittel, A. I., Malmstrom, R. R., and Cottrell, M. T. (2007). Standing stocks and activity of Archaea and Bacteria in the western Arctic Ocean. Limnol. Oceanogr. 52, 495–507. doi:10.4319/lo.2007.52.2.0495.

Korhonen, M., Rudels, B., Marnela, M., Wisotzki, A., and Zhao, J. (2013). Time and space variability of freshwater content, heat content and seasonal ice melt in the Arctic Ocean from 1991 to 2011. Ocean Sci. 9, 1015–1055. doi:10.5194/os-9-1015-2013.

Kraemer, S., Ramachandran, A., Colatriano, D., Lovejoy, C., and Walsh, D. A. (2020). Diversity and biogeography of SAR11 bacteria from the Arctic Ocean. ISME J. 14, 79–90. doi:10.1038/s41396-019-0499-4.

Kumar, M. S., Slud, E. V., Okrah, K., Hicks, S. C., Hannenhalli, S., and Bravo, H. C. (2017). Analysis and correction of compositional bias in sparse sequencing count data. bioRxiv 19, 799. doi:10.1101/142851.

Landry, Z., Swa, B. K., Herndl, G. J., Stepanauskas, R., and Giovannoni, S. J. (2017). SAR202 genomes from the dark ocean predict pathways for the oxidation of recalcitrant dissolved organic matter. MBio 8, e00413–17. doi:10.1128/mBio.00413-17.

Leu, E., Søreide, J. E., Hessen, D. O., Falk-Petersen, S., and Berge, J. (2011). Consequences of changing sea-ice cover for primary and secondary producers in the European Arctic shelf seas: Timing, quantity, and quality. Prog. Oceanogr. 90, 18–32. doi:10.1016/j.pocean.2011.02.004.

Luo, H., and Moran, M. A. (2014). Evolutionary Ecology of the Marine Roseobacter Clade. Microbiol. Mol. Biol. Rev. 78, 1–16. doi:10.1128/mmbr.88888-88.

Müller, O., Seuthe, L., Bratbak, G., and Paulsen, M. L. (2018a). Bacterial response to permafrost derived organic matter input in an Arctic Fjord. Front. Mar. Sci. 5. doi:10.3389/fmars.2018.00263.

Müller, O., Wilson, B., Paulsen, M. L., Ruminska, A., Armo, H. R., Bratbak, G., et al. (2018b). Spatiotemporal dynamics of ammonia-oxidizing Thaumarchaeota in Distinct Arctic water masses. Front. Microbiol. 9, 24. doi:10.3389/fmicb.2018.00024.

Mundy, C. J., Barber, D. G., and Michel, C. (2005). Variability of snow and ice thermal, physical and optical properties pertinent to sea ice algae biomass during spring. J. Mar. Syst. 58, 107–120. doi:10.1016/j.jmarsys.2005.07.003.

Nöthig, E.-M., Knüppel, N., and Lorenzen, C. (2018). Chlorophyll a measured on water bottle samples during POLARSTERN cruise PS99.2 (ARK-XXX/1.2). PANGAEA doi:10.1594/PANGAEA.887855.

Nöthig, E. M., Bracher, A., Engel, A., Metfies, K., Niehoff, B., Peeken, I., et al. (2015). Summertime plankton ecology in fram strait-a compilation of long-and short-term observations. Polar Res. 34, 23349. doi:10.3402/polar.v34.23349.

Owrid, G., Socal, G., Civitarese, G., Luchetta, A., Wiktor, J., Nöthig, E. M., et al. (2000). Spatial variability of phytoplankton, nutrients and new production estimates in the waters around Svalbard. Polar Res. 19, 155–171. doi:10.1111/j.1751-8369.2000.tb00340.x.

Peng, G., and Meier, W. N. (2018). Temporal and regional variability of Arctic sea-ice coverage from satellite data. Ann. Glaciol. 59, 191–200. doi:10.1017/aog.2017.32.

Pernthaler, A., Pernthaler, J., and Amann, R. (2002). Fluorescence in situ hybridization and catalyzed reporter deposition for the identification of marine bacteria. Appl. Environ. Microbiol. 68, 3094–3101. doi:10.1128/AEM.68.6.3094-3101.2002.

Perrette, M., Yool, A., Quartly, G. D., and Popova, E. E. (2011). Near-ubiquity of ice-edge blooms in the Arctic. Biogeosciences 8, 515–524. doi:10.5194/bg-8-515-2011.

Piontek, J., Sperling, M., Nöthig, E. M., and Engel, A. (2014). Regulation of bacterioplankton activity in Fram Strait (Arctic Ocean) during early summer: The role of organic matter supply and temperature. J. Mar. Syst. 132, 83–94. doi:10.1016/j.jmarsys.2014.01.003.

Piontek, J., Sperling, M., Nöthig, E. M., and Engel, A. (2015). Multiple environmental changes induce interactive effects on bacterial degradation activity in the arctic ocean. Limnol. Oceanogr. 60, 1392–1410. doi:10.1002/lno.10112.

Piwosz, K., Shabarova, T., Pernthaler, J., Posch, T., Šimek, K., Porcal, P., et al. (2020). Bacterial and Eukaryotic Small-Subunit Amplicon Data Do Not Provide a Quantitative Picture of Microbial Communities, but They Are Reliable in the Context of Ecological Interpretations. mSphere 5, 1–14. doi:10.1128/msphere.00052-20.

Polyakov, I. V., Pnyushkov, A. V., Alkire, M. B., Ashik, I. M., Baumann, T. M., Carmack, E. C., et al. (2017). Greater role for Atlantic inflows on sea-ice loss in the Eurasian Basin of the Arctic Ocean. Science (80-.). 356, 285–291. doi:10.1126/science.aai8204.

Quast, C., Pruesse, E., Yilmaz, P., Gerken, J., Schweer, T., Yarza, P., et al. (2013). The SILVA ribosomal RNA gene database project: Improved data processing and web-based tools. Nucleic Acids Res. 41, D590–D596. doi:10.1093/nar/gks1219.

Quero, G. M., Celussi, M., Relitti, F., Kovačević, V., Del Negro, P., and Luna, G. M. (2020). Inorganic and Organic Carbon Uptake Processes and Their Connection to Microbial Diversity in Meso- and Bathypelagic Arctic Waters (Eastern Fram Strait). Microb. Ecol. 79, 823–839. doi:10.1007/s00248-019-01451-2.

Redfield, A. C. (1963). “The influence of organisms on the composition of seawater,” in The sea (Wiley-Interscience), 26–77.

Rudels, B., Schauer, U., Björk, G., Korhonen, M., Pisarev, S., Rabe, B., et al. (2012). Observations of water masses and circulation in the Eurasian Basin of the Arctic Ocean from the 1990s to the late 2000s. Ocean Sci. Discuss. 9, 2695–2747. doi:10.5194/osd-9-2695-2012.

Salazar, G., Cornejo-Castillo, F. M., Benítez-Barrios, V., Fraile-Nuez, E., Álvarez-Salgado, X. A., Duarte, C. M., et al. (2016). Global diversity and biogeography of deep-sea pelagic prokaryotes. ISME J. 10, 596–608. doi:10.1038/ismej.2015.137.

Saw, J. H. W., Nunoura, T., Hirai, M., Takaki, Y., Parsons, R., Michelsen, M., et al. (2019). Pangenomics reveal diversification of enzyme families and niche specialization in globally abundant SAR202 bacteria. bioRxiv 11. doi:10.1101/692848.

Schattenhofer, M., Fuchs, B. M., Amann, R., Zubkov, M. V., Tarran, G. A., and Pernthaler, J. (2009). Latitudinal distribution of prokaryotic picoplankton populations in the Atlantic Ocean. Environ. Microbiol. 11, 2078–2093. doi:10.1111/j.1462-2920.2009.01929.x.

Schröder, M., and Wisotzki, A. (2014). Physical oceanography measured on water bottle samples during POLARSTERN cruise PS82 (ANT-XXIX/9). doi:10.1594/PANGAEA.871952.

Selje, N., Simon, M., and Brinkhoff, T. (2004). A newly discovered Roseobacter cluster in temperate and polar oceans. Nature 427, 445–448. doi:10.1038/nature02272.

Sheik, C. S., Jain, S., and Dick, G. J. (2014). Metabolic flexibility of enigmatic SAR324 revealed through metagenomics and metatranscriptomics. Environ. Microbiol. 16, 304–317. doi:10.1111/1462-2920.12165.

Soltwedel, T., Bauerfeind, E., Bergmann, M., Bracher, A., Budaeva, N., Busch, K., et al. (2016). Natural variability or anthropogenically-induced variation? Insights from 15 years of multidisciplinary observations at the arctic marine LTER site HAUSGARTEN. Ecol. Indic. 65, 89–102. doi:10.1016/j.ecolind.2015.10.001.

Soltwedel, T., Bauerfeind, E., Bergmann, M., Budaeva, N., Hoste, E., Jaeckisch, N., et al. (2005). Hausgarten: Multidisciplinary investigations at a Deep-Sea, long-term observatory in the Arctic Ocean. Oceanography 18, 46–61. doi:10.5670/oceanog.2005.24.

Sperling, M., Giebel, H. A., Rink, B., Grayek, S., Staneva, J., Stanev, E., et al. (2012). Differential effects of hydrographic and biogeochemical properties on the SAR11 clade and Roseobacter RCA cluster in the North Sea. Aquat. Microb. Ecol. 67, 25–34. doi:10.3354/ame01580.

Sun, L., Perlwitz, J., and Hoerling, M. (2016). What caused the recent “Warm Arctic, Cold Continents” trend pattern in winter temperatures? Geophys. Res. Lett. 43, 5345–5352. doi:10.1002/2016GL069024.

Swan, B. K., Martinez-Garcia, M., Preston, C. M., Sczyrba, A., Woyke, T., Lamy, D., et al. (2011). Potential for chemolithoautotrophy among ubiquitous bacteria lineages in the dark ocean. Science (80-.). 333, 1296–1300. doi:10.1126/science.1203690.

Tada, Y., Makabe, R., Kasamatsu-Takazawa, N., Taniguchi, A., and Hamasaki, K. (2013). Growth and distribution patterns of Roseobacter/Rhodobacter, SAR11, and Bacteroidetes lineages in the Southern Ocean. Polar Biol. 36, 691–704. doi:10.1007/s00300-013-1294-8.

Tada, Y., Taniguchi, A., Nagao, I., Miki, T., Uematsu, M., Tsuda, A., et al. (2011). Differing growth responses of major phylogenetic groups of marine bacteria to natural phytoplankton blooms in the Western North Pacific Ocean. Appl. Environ. Microbiol. 77, 4055–4065. doi:10.1128/AEM.02952-10.

Teeling, H., Fuchs, B. M., Becher, D., Klockow, C., Gardebrecht, A., Bennke, C. M., et al. (2012). Substrate-controlled succession of marine bacterioplankton populations induced by a phytoplankton bloom. Science (80-.). 336, 608–611. doi:10.1126/science.1218344.

Teeling, H., Fuchs, B. M., Bennke, C. M., Krüger, K., Chafee, M., Kappelmann, L., et al. (2016). Recurring patterns in bacterioplankton dynamics during coastal spring algae blooms. Elife 5, e11888. doi:10.7554/eLife.11888.

Teira, E., Lebaron, P., Van Aken, H., and Herndl, G. J. (2006). Distribution and activity of Bacteria and Archaea in the deep water masses of the North Atlantic. Limnol. Oceanogr. 51, 2131–2144. doi:10.4319/lo.2006.51.5.2131.

Teira, E., Reinthaler, T., Pernthaler, A., Pernthaler, J., and Herndl, G. J. (2004). Combining catalyzed reporter deposition-fluorescence in situ hybridization and microautoradiography to detect substrate utilization by bacteria and archaea in the deep ocean. Appl. Environ. Microbiol. 70, 4411–4414. doi:10.1128/AEM.70.7.4411-4414.2004.

Varela, M. M., Van Aken, H. M., and Herndl, G. J. (2008). Abundance and activity of Chloroflexi-type SAR202 bacterioplankton in the meso- and bathypelagic waters of the (sub)tropical Atlantic. Environ. Microbiol. 10, 1903–1911. doi:10.1111/j.1462-2920.2008.01627.x.

Vernet, M., Richardson, T. L., Metfies, K., Eva-Maria Nöthig, and Peeken, I. (2017). Models of plankton community changes during a warm water anomaly in Arctic waters show altered trophic pathways with minimal changes in carbon export. Front. Mar. Sci. 4, 160. doi:10.3389/fmars.2017.00160.

von Appen, W. J., Schauer, U., Somavilla, R., Bauerfeind, E., and Beszczynska-Möller, A. (2015). Exchange of warming deep waters across Fram Strait. Deep. Res. Part I Oceanogr. Res. Pap. 103, 86–100. doi:10.1016/j.dsr.2015.06.003.

Walczowski, W., Beszczynska-Möller, A., Wieczorek, P., Merchel, M., and Grynczel, A. (2017). Oceanographic observations in the Nordic Sea and Fram Strait in 2016 under the IO PAN long-term monitoring program AREX. Oceanologia 59, 187–194. doi:10.1016/j.oceano.2016.12.003.

Wassmann, P., and Reigstad, M. (2011). Future Arctic Ocean seasonal ice zones and implications for pelagic-benthic coupling. Oceanography 24, 220–231. doi:10.5670/oceanog.2011.74.

Wekerle, C., Wang, Q., von Appen, W. J., Danilov, S., Schourup-Kristensen, V., and Jung, T. (2017). Eddy-Resolving Simulation of the Atlantic Water Circulation in the Fram Strait With Focus on the Seasonal Cycle. J. Geophys. Res. Ocean. 122, 8385–8405. doi:10.1002/2017JC012974.

Welch, D. B. M., and Huse, S. M. (2011). Microbial Diversity in the Deep Sea and the Underexplored “Rare Biosphere.” Handb. Mol. Microb. Ecol. II Metagenomics Differ. Habitats 103, 243–252. doi:10.1002/9781118010549.ch24.

Wells, L. E., Cordray, M., Bowerman, S., Miller, L. A., Vincent, W. F., and Deming, J. W. (2006). Archaea in particle-rich waters of the Beaufort Shelf and Franklin Bay, Canadian Arctic: Clues to an allochthonous origin? Limnol. Oceanogr. 51, 47–59. doi:10.4319/lo.2006.51.1.0047.

Wells, L. E., and Deming, J. W. (2003). Abundance of bacteria, the Cytophaga-Flavobacterium cluster and Archaea in cold oligotrophic waters and nepheloid layers of the Northwest Passage, Canadian archipelago. Aquat. Microb. Ecol. 31, 19–31. doi:10.3354/ame031019.

Wickham, H. (2016). “Getting Started with ggplot2,” in ggplot2 (Springer), 11–31. doi:10.1007/978-3-319-24277-4_2.

Wickham, H., Averick, M., Bryan, J., Chang, W., McGowan, L., François, R., et al. (2019). Welcome to the Tidyverse. J. Open Source Softw. 4, 1686. doi:10.21105/joss.01686.

Wilson, B., Müller, O., Nordmann, E. L., Seuthe, L., Bratbak, G., and Øvreås, L. (2017). Changes in marine prokaryote composition with season and depth over an Arctic polar year. Front. Mar. Sci. 4, 95. doi:10.3389/fmars.2017.00095.

Wilson, C., and Wallace, D. W. R. (1990). Using the nutrient ratio NO/PO as a tracer of continental shelf waters in the central Arctic Ocean. J. Geophys. Res. 95, 22193. doi:10.1029/jc095ic12p22193.

Zeder, M., Ellrott, A., and Amann, R. (2011). Automated sample area definition for high-throughput microscopy. Cytom. Part A 79 A, 306–310. doi:10.1002/cyto.a.21034.

