## Supplementary information for "Spatial distribution of Arctic bacterioplankton abundance is linked to distinct water masses and summertime phytoplankton bloom dynamics (Fram Strait, 79°N)"

**Table S1.** Environmental parameters measured at the different stations. Chl. a: Chlorophyll a. EGC: East Greenland current region, N: North region, WSC: West Spitsbergen Current region. SUR: surface mixed water, EPI: epipelagic, MES: mesopelagic, and BAT: bathypelagic zone. Lat: latitude. Lon: longitude. Temp: Temperature. Sal: Salinity.

| Region | Station | PANGAEA Event ID | Water layer | Lat (°N) | Lon (°E) | Depth (m) | Temp (°C) | Sal | Chl *a* (µg $L^{-1}$) | $NO_3$ (µmol $L^{-1}$) | $\Delta NO_3$ (µmol $L^{-1}$) | $PO_4$ (µmol $L^{-1}$) | $\Delta PO_4$ (µmol $L^{-1}$) | N:P | $NH_4$ (µmol $L^{-1}$) | $\Delta NH_4$ (µmol $L^{-1}$) | $SiO_3$ (µmol $L^{-1}$) | $\Delta SiO_3$ (µmol $L^{-1}$) |
|---|---|---|---|---|---|---|---|---|---|---|---|---|---|---|---|---|---|---|
| EGC | EG1 | PS99/051-2 | SUR | 78.99 | -5.42 | 13 | -1.51 | 33.17 | 1.66 | 3.75 ± 0.01 | 3.55 ± 0.01 | 0.46 ± 0.00 | 0.13 ± 0.00 | 8.15 | 0 ± 0.00 | 0.00 | 4.50 ± 0.00 | 0.14 ± 0.02 |
| EGC | EG1 | PS99/051-2 | EPI | 78.99 | -5.42 | 100 | -1.11 | 34.23 | | 8.44 ± 0.04 | | 0.63 ± 0.00 | | 13.40 | 0 ± 0.01 | | 4.04 ± 0.00 | |
| EGC | EG1 | PS99/051-2 | MES | 78.99 | -5.42 | 971 | -0.14 | 34.89 | | 12.62 ± 0.03 | | 0.85 ± 0.00 | | 14.85 | 0 ± 0.03 | | 7.00 ± 0.04 | |
| EGC | EG4 | PS99/048-11 | SUR | 78.82 | -2.73 | 24 | -1.17 | 34.48 | 1.52 | 4.92 ± 0.02 | 6.34 ± 0.01 | 0.50 ± 0.00 | 0.35 ± 0.00 | 9.84 | 0 ± 0.02 | 0.08 ± 0.01 | 2.44 ± 0.14 | 2.38 ± 0.34 |
| EGC | EG4 | PS99/048-11 | EPI | 78.82 | -2.73 | 100 | 3.86 | 35.05 | | 10.97 ± 0.07 | | 0.81 ± 0.00 | | 13.54 | 0 ± 0.02 | | 4.83 ± 0.17 | |
| EGC | EG4 | PS99/048-1 | MES | 78.82 | -2.73 | 1000 | -0.24 | 34.91 | | 13.77 ± 0.04 | | 0.94 ± 0.02 | | 14.65 | 0 ± 0.01 | | 7.10 ± 0.01 | |
| EGC | EG4 | PS99/048-1 | BAT | 78.82 | -2.73 | 2527 | -0.76 | 34.91 | | 14.97 ± 0.09 | | 1.06 ± 0.01 | | 14.12 | 0 ± 0.01 | | 12.07 ± 0.19 | |
| N | N3 | PS99/054-1 | SUR | 79.58 | 5.17 | 34 | 3.28 | 34.40 | 0.95 | 8.09 ± 0.06 | 5.23 ± 0.01 | 0.72 ± 0.00 | 0.31 ± 0.00 | 11.24 | 1.29 ± 0.00 | 0.00 | 3.19 ± 0.05 | 1.02 ± 0.03 |
| N | N3 | PS99/054-1 | EPI | 79.58 | 5.17 | 100 | 4.25 | 35.10 | | 10.11 ± 0.04 | | 0.78 ± 0.00 | | 12.96 | 0.47 ± 0.01 | | 3.29 ± 0.04 | |
| N | N3 | PS99/054-1 | MES | 79.58 | 5.17 | 1000 | -0.33 | 34.10 | | 13.51 ± 0.02 | | 1.05 ± 0.00 | | 12.87 | 0 ± 0.00 | | 7.25 ± 0.00 | |
| N | N3 | PS99/054-1 | BAT | 79.58 | 5.17 | 2500 | -0.72 | 34.92 | | 14.81 ± 0.05 | | 1.09 ± 0.00 | | 13.59 | 0 ± 0.00 | | 10.74 ± 0.03 | |
| N | N4 | PS99/055-1 | SUR | 79.74 | 4.51 | 22 | 2.66 | 33.99 | 2.21 | 3.33 ± 0.01 | 6.80 ± 0.01 | 0.53 ± 0.01 | 0.49 ± 0.01 | 6.28 | 0.29 ± 0.03 | 0.00 | 1.65 ± 0.00 | 2.06 ± 0.01 |
| N | N4 | PS99/055-1 | EPI | 79.74 | 4.51 | 100 | 3.94 | 35.08 | | 10.64 ± 0.00 | | 1.00 ± 0.03 | | 10.64 | 0.42 ± 0.01 | | 3.90 ± 0.00 | |
| N | N4 | PS99/055-7 | MES | 79.74 | 4.51 | 1000 | -0.41 | 34.91 | | 13.96 ± 0.08 | | 0.92 ± 0.00 | | 15.17 | 0 ± 0.01 | | 9.07 ± 0.19 | |
| N | N4 | PS99/055-7 | BAT | 79.74 | 4.51 | 2500 | -0.74 | 34.92 | | 14.47 ± 0.02 | | 0.86 ± 0.00 | | 16.83 | 0 ± 0.00 | | 11.24 ± 0.14 | |
| N | N5 | PS99/053-2 | SUR | 79.92 | 3.06 | 19 | 0.75 | 33.59 | 7.40 | 0.97 ± 0.01 | 8.21 ± 0.01 | 0.51 ± 0.04 | 0.66 ± 0.02 | 1.90 | 0 ± 0.04 | 1.56 ± 0.01 | 0.71 ± 0.01 | 2.34 ± 0.04 |
| N | N5 | PS99/053-2 | EPI | 79.92 | 3.06 | 100 | 4.27 | 35.10 | | 10.05 ± 0.05 | | 1.10 ± 0.06 | | 9.14 | 0.42 ± 0.01 | | 3.22 0.08 | |
| N | N5 | PS99/053-2 | MES | 79.92 | 3.06 | 1000 | -0.23 | 34.91 | | 13.04 ± 0.12 | | 1.34 ± 0.01 | | 9.73 | 0 ± 0.00 | | 6.50 ± 0.02 | |
| N | N5 | PS99/053-2 | BAT | 79.92 | 3.06 | 2427 | -0.74 | 34.92 | | 14.29 ± 0.04 | | 1.59 ± 0.00 | | 8.99 | 0 ± 0.01 | | 10.79 ± 0.01 | |
| WSC | HG1 | PS99/066-2 | SUR | 79.14 | 6.09 | 17 | 6.27 | 35.33 | 3.42 | | | | | | | | | |
| WSC | HG1 | PS99/066-5 | EPI | 79.14 | 6.09 | 100 | 4.38 | 35.09 | | | | | | | | | | |

| | | | | | | | | | | | | | | | | | | |
|---|---|---|---|---|---|---|---|---|---|---|---|---|---|---|---|---|---|---|
| WSC | HG1 | PS99/066-5 | MES | 79.14 | 6.09 | 500 | 1.46 | 34.95 | | | | | | | | | | |
| WSC | HG1 | PS99/066-5 | BAT | 79.14 | 6.09 | 1253 | -0.81 | 34.91 | | | | | | | | | | |
| WSC | HG2 | PS99/057-1 | SUR | 79.13 | 4.91 | 22 | 2.30 | 34.90 | 2.23 | 6.15 ± 0.01 | 5.30 ± 0.01 | 0.89 ± 0.00 | 0.55 ± 0.02 | 6.91 | 0.94 ± 0.00 | 0.00 | 2.74 ± 0.02 | 1.36 ± 0.05 |
| WSC | HG2 | PS99/057-1 | EPI | 79.13 | 4.91 | 100 | 3.60 | 35.04 | | 10.84 ± 0.01 | | 1.30 ± 0.01 | | 8.34 | 0 ± 0.01 | | 3.88 ± 0.11 | |
| WSC | HG2 | PS99/057-1 | MES | 79.13 | 4.91 | 1000 | -0.58 | 34.91 | | 14.15 ± 0.09 | | 1.69 ± 0.01 | | 8.37 | 0 ± 0.01 | | 9.33 ± 0.06 | |
| WSC | HG2 | PS99/057-1 | BAT | 79.13 | 4.91 | 1492 | -0.81 | 34.91 | | 14.95 ± 0.02 | | 1.78 ± 0.07 | | 8.40 | 0 ± 0.00 | | 12.35 ± 0.02 | |
| WSC | HG4 | PS99/042-11 | SUR | 79.07 | 4.19 | 28 | 0.41 | 34.44 | 3.54 | 5.79 ± 0.04 | 6.42 ± 0.04 | 0.66 ± 0.02 | 0.41 ± 0.02 | 8.77 | 0.56 ± 0.01 | 0.17 ± 0.01 | 2.66 ± 0.02 | 2.11 ± 0.03 |
| WSC | HG4 | PS99/042-11 | EPI | 79.07 | 4.19 | 100 | 3.51 | 35.04 | | 10.88 ± 0.07 | | 0.87 ± 0.06 | | 12.51 | 0 ± 0.01 | | 4.16 ± 0.12 | |
| WSC | HG4 | PS99/042-1 | MES | 79.06 | 4.19 | 1000 | -0.35 | 34.91 | | 13.71 ± 0.04 | | 0.96 ± 0.01 | | 14.28 | 0 ± 0.01 | | 7.70 ± 0.02 | |
| WSC | HG4 | PS99/042-1 | BAT | 79.06 | 4.19 | 2462 | -0.73 | 34.92 | | 14.67 ± 0.05 | | 1.01 ± 0.00 | | 14.52 | 0 ± 0.00 | | 11.59 ± 0.03 | |
| WSC | HG5 | PS99/044-1 | SUR | 79.07 | 3.66 | 25 | 1.74 | 32.363 | 4.24 | 5.1 ± 0.03 | 7.08 ± 0.02 | 0.53 ± 0.00 | 0.37 ± 0.00 | 9.62 | 0.50 ± 0.02 | 0.00 | 3.19 ± 0.01 | 1.96 ± 0.06 |
| WSC | HG5 | PS99/044-1 | EPI | 79.07 | 3.66 | 100 | 3.94 | 35.089 | | 11.26 ± 0.03 | | 0.80 ± 0.00 | | 14.08 | 0 ± 0.01 | | 4.75 ± 0.03 | |
| WSC | HG5 | PS99/044-1 | MES | 79.07 | 3.66 | 2600 | -0.73 | 34.924 | | | | | | | | | | |
| WSC | HG5 | PS99/044-1 | BAT | 79.07 | 3.66 | 3038 | -0.70 | 34.925 | | | | | | | | | | |
| WSC | HG7 | PS99/046-1 | SUR | 79.05 | 3.53 | 35 | 3.73 | 33.957 | 4.46 | 5.24 ± 0.01 | 2.61 ± 0.05 | 0.51 ± 0.00 | 0.09 ± 0.00 | 10.27 | 0.48 ± 0.00 | 0.00 | 2.84 ± 0.20 | 1.68 ± 0.08 |
| WSC | HG7 | PS99/046-1 | EPI | 79.05 | 3.53 | 100 | 3.35 | 35.016 | | 10.35 ± 0.04 | | 0.74 ± 0.00 | | 13.99 | 0.09 ± 0.00 | | 4.21 ± 0.02 | |
| WSC | HG7 | PS99/046-1 | MES | 79.05 | 3.53 | 1005 | -0.28 | 34.908 | | | | | | | | | | |
| WSC | HG7 | PS99/046-1 | BAT | 79.05 | 3.53 | 3772 | -0.63 | 34.924 | | | | | | | | | | |
| WSC | HG9 | PS99/059-2 | SUR | 79.13 | 2.84 | 24 | -1.24 | 35.089 | 1.90 | 1.74 ± 0.04 | 6.58 ± 0.01 | 0.67 ± 0.00 | 0.58 ± 0.13 | 2.60 | 0 ± 0.02 | 0.73 ± 0.21 | 1.72 ± 0.02 | 1.96 ± 0.01 |
| WSC | HG9 | PS99/059-2 | EPI | 79.13 | 2.84 | 100 | 3.92 | 35.047 | | 10.97 ± 0.01 | | 1.12 ± 0.05 | | 9.79 | 0.47 ± 0.28 | | 3.70 ± 0.03 | |
| WSC | HG9 | PS99/059-2 | MES | 79.13 | 2.84 | 1000 | -0.19 | 34.897 | | 13.59 ± 0.02 | | 1.17 ± 0.17 | | 11.62 | 0.00 ± 0.17 | | 6.82 ± 0.01 | |
| WSC | HG9 | PS99/059-2 | BAT | 79.13 | 2.84 | 2499 | -0.72 | 34.919 | | 15.23 ± 0.01 | | 1.27 ± 0.03 | | 11.99 | 0.00 ± 0.00 | | 11.11 ± 0.03 | |

**Table S2**. Pearson's correlation coefficient (*r*) tests between environmental parameters, diatoms and *Phaeocystis* spp. counts and cell abundances of the investigated taxonomic groups in surface waters of the Fram Strait. Combinations that show significant correlation are marked with grey shadow. The number of data pairs was 10. *Bacteria* (EUB), *Archaea* (ARCH), *Alteromonadaceae/Colwelliaceae/Pseudoalteromonadaceae* (ALT). *Bacteroidetes* (BACT). *Chloroflexi* (CFX). *Thaumarchaeota* (THA). *Deltaproteobacteria* (DELTA). *Gammaproteobacteria* (GAM). *Opitutales* (OPI). *Polaribacter* (POL). *Rhodobacteraceae* (ROS). SAR11. SAR202. SAR324. SAR406. *Verrucomicrobiales* (VER).

| **Taxa** | **Temperature** | | **Salinity** | | **Chlorophyll *a* conc.** | | **$\Delta NO_3$** | | **$\Delta PO_4$** | | **$\Delta SO_3$** | | **Diatoms** | | ***Phaeocystis.spp*** | |
|---|---|---|---|---|---|---|---|---|---|---|---|---|---|---|---|---|
| | ***r*** | ***p-value*** | ***r*** | ***p-value*** | ***r*** | ***p-value*** | ***r*** | ***p-value*** | ***r*** | ***p-value*** | ***r*** | ***p-value*** | ***r*** | ***p-value*** | ***r*** | ***p-value*** |
| ALT | 0.03 | 0.93 | 0.28 | 0.41 | -0.28 | 0.40 | -0.12 | 0.74 | 0.27 | 0.44 | -0.23 | 0.519 | 0.49 | 0.40 | -0.84 | 0.08 |
| ARCH | 0.47 | 0.14 | 0.36 | 0.28 | -0.50 | 0.12 | -0.31 | 0.39 | -0.21 | 0.56 | -0.63 | 0.05 | 0.87 | 0.06 | -0.15 | 0.81 |
| BACT | 0.41 | 0.21 | 0.52 | 0.10 | -0.25 | 0.46 | 0.14 | 0.70 | 0.56 | 0.09 | 0.01 | 0.98 | 0.40 | 0.50 | -0.36 | 0.55 |
| CFX | 0.12 | 0.72 | 0.40 | 0.23 | -0.25 | 0.45 | 0.07 | 0.85 | 0.45 | 0.19 | -0.12 | 0.74 | -0.22 | 0.72 | -0.38 | 0.53 |
| THA | 0.35 | 0.30 | 0.59 | 0.06 | -0.30 | 0.37 | -0.1 | 0.79 | -0.09 | 0.80 | -0.05 | 0.90 | -0.19 | 0.76 | 0.78 | 0.12 |
| DELTA | -0.4 | 0.23 | 0.05 | 0.89 | -0.43 | 0.18 | 0.04 | 0.91 | -0.15 | 0.68 | -0.60 | 0.07 | 0.38 | 0.53 | **-0.89** | **0.04** |
| EUB | 0.60 | 0.05 | 0.50 | 0.12 | 0.22 | 0.52 | 0.17 | 0.64 | 0.59 | 0.07 | 0.14 | 0.69 | -0.32 | 0.60 | 0.19 | 0.76 |
| GAM | 0.14 | 0.69 | 0.04 | 0.91 | 0.50 | 0.12 | 0.42 | 0.22 | **0.74** | **0.01** | 0.41 | 0.25 | -0.34 | 0.57 | 0.08 | 0.90 |
| OPI | **0.61** | **0.05** | 0.38 | 0.25 | -0.02 | 0.94 | 0.06 | 0.87 | 0.49 | 0.15 | -0.02 | 0.96 | -0.01 | 0.98 | 0.11 | 0.85 |
| POL | -0.10 | 0.78 | 0.00 | 0.99 | 0.59 | 0.05 | 0.33 | 0.36 | 0.54 | 0.12 | 0.25 | 0.48 | -0.30 | 0.62 | 0.03 | 0.96 |
| ROS | **0.78** | **0.00** | 0.49 | 0.13 | 0.06 | 0.87 | 0.26 | 0.47 | 0.54 | 0.12 | 0.09 | 0.80 | -0.13 | 0.84 | 0.32 | 0.60 |
| SAR11 | **0.79** | **0.00** | **0.61** | **0.04** | -0.09 | 0.78 | -0.15 | 0.69 | 0.30 | 0.40 | -0.08 | 0.82 | 0.02 | 0.97 | 0.43 | 0.47 |
| SAR202 | 0.16 | 0.64 | 0.29 | 0.39 | -0.51 | 0.11 | 0.23 | 0.51 | 0.33 | 0.36 | -0.37 | 0.29 | 0.82 | 0.09 | -0.33 | 0.59 |
| SAR324 | 0.14 | 0.69 | **0.64** | **0.03** | -0.39 | 0.24 | 0.21 | 0.56 | 0.22 | 0.55 | 0.03 | 0.93 | 0.02 | 0.97 | 0.77 | 0.13 |
| SAR406 | 0.04 | 0.92 | 0.26 | 0.44 | -0.33 | 0.32 | 0.18 | 0.62 | 0.18 | 0.63 | -0.29 | 0.41 | -0.58 | 0.30 | -0.18 | 0.77 |
| VER | **0.68** | **0.02** | 0.06 | 0.86 | 0.20 | 0.55 | -0.18 | 0.61 | 0.20 | 0.59 | 0.10 | 0.78 | -0.10 | 0.87 | 0.17 | 0.78 |

**Table S3**. Average bacterioplankton cell abundances along the water column of ice-covered and ice-free regions of the Fram Strait. The DAPI counts represent total bacterioplankton cell abundances. The proportions (%) of *Archaea* (ARCH) and *Bacteria* (EUB) were calculated based on the total bacterioplankton cell abundances. Sample size 'n' represents the number of counted fields of view. Standard error was not calculated for samples of the EGC located in the bathypelagic zone due to one station located at this depth in the region (NA). EGC: the ice-covered East Greenland Current stations, N: the marginal ice northern stations, WSC: the ice-free West Spitsbergen Current stations. All values are represented in $10^5$ cells $mL^{-1}$.

| **Region** | **Water layer** | **DAPI** | ***Archaea*** | **%** | ***n*** | ***Bacteria*** | **%** | ***n*** |
|---|---|---|---|---|---|---|---|---|
| EGC | Surface | 3.4 ± 0.2 | 0.2 ± 0.0 | 8 | 56 | 2.2 ± 0.2 | 60 | 52 |
| EGC | Epipelagic | 3.5 ± 1.2 | 0.4 ± 0.2 | 14 | 57 | 2.1 ± 1.3 | 55 | 48 |
| EGC | Mesopelagic | 0.7 ± 0.3 | 0.1 ± 0.1 | 17 | 51 | 0.2 ± 0.1 | 40 | 44 |
| EGC | Bathypelagic | 0.6 ± NA | 0.02 ± NA | 12 | 32 | 0.1 ± NA | 16 | 32 |
| N | Surface | 17.1 ± 0.7 | 0.3 ± 0.1 | 2 | 90 | 10.7 ± 0.7 | 62 | 81 |
| N | Epipelagic | 7.9 ± 1.2 | 0.6 ± 0.0 | 9 | 90 | 3.2 ± 0.7 | 40 | 73 |
| N | Mesopelagic | 0.9 ± 0.1 | 0.1 ± 0.0 | 17 | 77 | 0.3 ± 0.0 | 37 | 101 |
| N | Bathypelagic | 0.4 ± 0.0 | 0.1 ± 0.0 | 17 | 63 | 0.1 ± 0.0 | 37 | 85 |
| WSC | Surface | 15.0 ± 3.6 | 0.2 ± 0.0 | 1 | 146 | 8.1 ± 1.8 | 59 | 150 |
| WSC | Epipelagic | 6.2 ± 0.7 | 0.6 ± 0.1 | 12 | 217 | 2.2 ± 0.3 | 36 | 175 |
| WSC | Mesopelagic | 0.8 ± 0.1 | 0.1 ± 0.0 | 13 | 152 | 0.3 ± 0.1 | 34 | 166 |
| WSC | Bathypelagic | 0.5 ± 0.1 | 0.1 ± 0.0 | 15 | 198 | 0.2 ± 0.0 | 33 | 201 |

**Table S4.** Cell abundances of all taxonomic groups ($10^5$ cells $mL^{-1}$), and their proportions (%) towards bacterioplankton in the different Fram Strait regions: *Alteromonadaceae/Colwelliaceae/Pseudoalteromonadaceae* (ALT), *Bacteroidetes* (BACT), *Chloroflexi* (CFX), *Thaumarchaeota* (THA), *Deltaproteobacteria* (DELTA), *Gammaproteobacteria* (GAM), *Opitutales* (OPI), *Polaribacter* (POL), *Rhodobacteraceae* (ROS), SAR11, SAR202, SAR324, SAR406 and *Verrucomicrobiales* (VER). n: sample size (Fields of View). EGC: East Greenland current region, N: North region, WSC: West Spitsbergen Current region. SUR: surface mixed water, EPI: epipelagic, MES: mesopelagic, and BAT: bathypelagic zone.

| **Region** | **Water layer** | **ALT** | **%** | ***n*** | **BACT** | **%** | ***n*** | **CFX** | **%** | ***n*** | **SAR202** | **%** | ***n*** | **THA** | **%** | ***n*** | **DELTA** | **%** | ***n*** | **SAR406** | **%** | ***n*** |
|---|---|---|---|---|---|---|---|---|---|---|---|---|---|---|---|---|---|---|---|---|---|---|
| EGC | SUR | 0.2 ± 0.1 | 7 | 64 | 0.6 ± 0.4 | 18 | 66 | 0.06 ± 0.01 | 2 | 48 | 0.06 ± 0.01 | 2 | 44 | 0.1 ± 0.0 | 3 | 34 | 0.2 ± 0.1 | 7 | 57 | 0.03 ± 0.0 | 1 | 13 |
| EGC | EPI | 0.1 ± 0.0 | 2 | 61 | 0.3 ± 0.2 | 8 | 93 | 0.06 ± 0.01 | 2 | 51 | 0.06 ± 0.01 | 2 | 53 | 0.1 ± 0.02 | 4 | 75 | 0.1 ± 0.0 | 4 | 67 | 0.04 ± 0.01 | 2 | 6 |
| EGC | MES | 0.02 ± 0.0 | 3 | 53 | 0.01 ± 0.0 | 2 | 23 | 0.02 ± 0.0 | 5 | 57 | 0.02 ± 0.0 | 4 | 55 | 0.01 ± 0.0 | 2 | 27 | 0.03 ± 0.01 | 5 | 45 | 0.01 ± 0.0 | 1 | 11 |
| EGC | BAT | 0.01 ± NA | 2 | 11 | 0.01 ± NA | 1 | 25 | 0.02 ± NA | 4 | 18 | 0.02 ± NA | 5 | 26 | 0.01 ± NA | 2 | 14 | 0.02 ± NA | 3 | 33 | 0.01 ± NA | 3 | 6 |
| N | SUR | 0.1 ± 0.0 | 1 | 68 | 2.1 ± 0.8 | 12 | 85 | 0.2 ± 0.1 | 1 | 70 | 0.07 ± 0.01 | 1 | 57 | 0.1 ± 0.0 | 1 | 65 | 0.2 ± 0.1 | 1 | 86 | 0.06 ± 0.0 | 1 | 47 |
| N | EPI | 0.2 ± 0.1 | 2 | 87 | 0.4 ± 0.1 | 6 | 102 | 0.17 ± 0.1 | 2 | 107 | 0.05 ± 0.01 | 1 | 83 | 0.3 ± 0.0 | 4 | 122 | 0.2 ± 0.0 | 3 | 94 | 0.04 ± 0.0 | 1 | 28 |
| N | MES | 0.02 ± 0.0 | 2 | 73 | 0.02 ± 0.0 | 2 | 51 | 0.03 ± 0.0 | 3 | 116 | 0.03 ± 0.0 | 3 | 107 | 0.01 ± 0.0 | 1 | 28 | 0.04 ± 0.0 | 5 | 99 | 0.01 ± 0.0 | 1 | 18 |
| N | BAT | 0.02 ± 0.0 | 5 | 53 | 0.01 ± 0.0 | 3 | 26 | 0.02 ± 0.0 | 6 | 115 | 0.03 ± 0.0 | 8 | 120 | 0.01 ± 0.0 | 4 | 39 | 0.03 ± 0.0 | 8 | 123 | 0.01 ± 0.0 | 4 | 37 |
| WSC | SUR | 0.3 ± 0.1 | 2 | 186 | 2.1 ± 1.0 | 16 | 163 | 0.3 ± 0.2 | 3 | 127 | 0.05 ± 0.01 | 1 | 87 | 0.1 ± 0.0 | 1 | 178 | 0.1 ± 0.0 | 1 | 121 | 0.04 ± 0.0 | 1 | 71 |
| WSC | EPI | 0.1 ± 0.0 | 1 | 186 | 0.2 ± 0.0 | 4 | 195 | 0.1 ± 0.03 | 2 | 183 | 0.05 ± 0.01 | 1 | 196 | 0.3 ± 0.1 | 5 | 246 | 0.2 ± 0.1 | 4 | 196 | 0.03 ± 0.01 | 1 | 64 |
| WSC | MES | 0.02± 0.0 | 3 | 90 | 0.01 ± 0.0 | 2 | 71 | 0.03 ± 0.0 | 3 | 217 | 0.03 ± 0.0 | 5 | 215 | 0.02 ± 0.0 | 3 | 86 | 0.1± 0.01 | 6 | 139 | 0.01 ± 0.0 | 2 | 37 |
| WSC | BAT | 0.02 ± 0.0 | 4 | 102 | 0.01 ± 0.0 | 3 | 50 | 0.02 ± 0.0 | 4 | 199 | 0.03 ± 0.0 | 7 | 210 | 0.02 ± 0.0 | 4 | 54 | 0.1 ± 0.02 | 9 | 130 | 0.01 ± 0.0 | 3 | 45 |
| **Region** | **Water layer** | **GAM** | **%** | ***n*** | **OPI** | **%** | ***n*** | **POL** | **%** | ***n*** | **ROS** | **%** | ***n*** | **11 SAR** | **%** | ***n*** | **324 SAR** | **%** | ***n*** | **VER** | **%** | ***n*** |
| EGC | SUR | 0.32 ± 0.2 | 125 | 67 | 0.1 ± 0.0 | 2 | 53 | 0.4 ± 0.3 | 11 | 62 | 0.2 ± 0.1 | 7 | 44 | 1.9 ± 0.7 | 50 | 53 | 0.1 ± 0.04 | 2 | 44 | 0.1 ± 0.0 | 3 | 41 |
| EGC | EPI | 0.2 ± 0.1 | 5 | 77 | 0.1 ± 0.0 | 2 | 70 | 0.1 ± 0.0 | 2 | 52 | 0.3 ± 0.2 | 9 | 60 | 1.0 ± 0.6 | 26 | 50 | 0.1 ± 0.1 | 3 | 39 | 0.1 ± 0.0 | 12 | 63 |
| EGC | MES | 0.01 ± 0.0 | 2 | 27 | 0.02 ± 0.0 | 3 | 40 | 0.01 ± 0.0 | 3 | 47 | 0.1 ± 0.0 | 11 | 75 | 0.1 ± 0.0 | 22 | 49 | 0.1 ± 0.0 | 6 | 60 | 0.01 ± 0.0 | 3 | 37 |
| EGC | BAT | 0.01 ± NA | 2 | 5 | 0.01 ± NA | 3 | 11 | 0.02 ± NA | 2 | 29 | 0.03 ± NA | 5 | 14 | 0.1 ± NA | 14 | 32 | 0.03 ± NA | 5 | 35 | 0.01 ± NA | 2 | 11 |

| | | | | | | | | | | | | | | | | | | | | | |
|---|---|---|---|---|---|---|---|---|---|---|---|---|---|---|---|---|---|---|---|---|---|
| N | SUR | 2.1 ± 0.6 | 128 | 78 | 0.9 ± 0.01 | 5 | 91 | 1.3 ± 0.6 | 8 | 85 | 0.6 ± 0.0 | 3 | 92 | 4.2 ± 1.0 | 24 | 81 | 0.07 ± 0.01 | 1 | 80 | 0.9 ± 0. 1 | 5 | 91 |
| N | EPI | 0.4 ± 0.1 | 5 | 109 | 0.2 ± 0.1 | 2 | 107 | 0.1 ± 0.0 | 2 | 98 | 0.2 ± 0.0 | 3 | 108 | 2.2 ± 0.3 | 29 | 73 | 0.07 ± 0.01 | 1 | 97 | 0.2 ± 0.1 | 2 | 98 |
| N | MES | 0.02 ± 0.0 | 2 | 93 | 0.02 ± 0.0 | 2 | 74 | 0.02± 0.0 | 2 | 70 | 0.1 ± 0.01 | 13 | 128 | 0.2 ± 0.0 | 20 | 101 | 0.06 ± 0.0 | 6 | 66 | 0.02 ± 0.0 | 2 | 73 |
| N | BAT | 0.02 ± 0.0 | 4 | 55 | 0.01 ± 0.0 | 5 | 52 | 0.02 ± 0.0 | 5 | 55 | 0.1 ± 0.0 | 20 | 122 | 0.1 ± 0.0 | 27 | 85 | 0.03 ± 0.0 | 8 | 54 | 0.01 ± 0.0 | 4 | 48 |
| WSC | SUR | 1.6 ± 0.3 | 136 | 189 | 0.8 ± 0.3 | 5 | 172 | 0.8 ± 0.2 | 7 | 162 | 0.7 ± 0.3 | 4 | 165 | 6.1 ± 2.0 | 38 | 150 | 0.07 ± 0.02 | 1 | 146 | 1.1 ± 0.0 | 5 | 171 |
| WSC | EPI | 0.2 ± 0.0 | 3 | 214 | 0.1 ± 0.0 | 2 | 238 | 0.1 ± 0.0 | 1 | 176 | 0.1 ± 0.0 | 1 | 223 | 1.9 ± 0.2 | 32 | 175 | 0.09 ± 0.01 | 2 | 141 | 0.1 ± 0.0 | 2 | 219 |
| WSC | MES | 0.01 ± 0.0 | 3 | 108 | 0.02± 0.0 | 2 | 110 | 0.02 ± 0.0 | 2 | 128 | 0.1 ± 0.01 | 7 | 195 | 0.2 ± 0.0 | 18 | 166 | 0.05 ± 0.0 | 5 | 168 | 0.02 ± 0.0 | 3 | 98 |
| WSC | BAT | 0.02 ± 0.0 | 3 | 105 | 0.02 ± 0.0 | 5 | 110 | 0.02 ± 0.0 | 5 | 129 | 0.04 ± 0.01 | 14 | 197 | 0.1 ± 0.0 | 21 | 201 | 0.05 ± 0.0 | 6 | 138 | 0.02 ± 0.0 | 4 | 74 |

**Table S5.** Specificities of rRNA-targeting oligonucleotide probes used during CARD-FISH for the quantification of pelagic microbial groups in the Fram Strait. FA: formamide concentration in the hybridization buffer.

| Probe name | Target group | Sequence (5’-3’) | FA (%) | Reference |
|---|---|---|---|---|
| EUB338 I | *Bacteria* | GCT GCC TCC CGT AGG AGT | 35 | (Amann et al., 1990) |
| EUB338 II | *Planctomycetales* | GCA GCC ACC CGT AGG TGT | 35 | (Daims et al., 1999) |
| EUB338 III | *Verrucomicrobiales* | GCT GCC ACC CGT AGG TGT | 35 | (Daims et al., 1999) |
| Non338 | nonsense probe | ACT CCT ACG GGA GGC AGC | 35 | (Wallner et al., 1993) |
| ARCH915 | *Archaea* | GTG CTC CCC CGC CAA TTC CT | 35 | (Amann et al., 1995) |
| CFX1223 | *Chloroflexi* | CCA TTG TAG CGT GTG TGT MG | 35 | (Björnsson et al., 2002) |
| GNSB941 | *Chloroflexi* | AAA CCA CAC GCT CCG CT | 35 | (Gich et al., 2001) |
| SAR202-312R | SAR202 clade | TGT CTC AGT CCC CCT CTG | 40 | (Morris et al., 2004) |
| PSA184 | *Alteromonadaceae, Colwelliaceae, Pseudoalteromonadaceae* | CCC CTT TGG TCC GTA GAC | 30 | (Eilers et al., 2000) |
| GAM42a | *Gammaproteobacteria* | GCC TTC CCA CAT CGT TT | 35 | (Manz et al., 1992) |
| BET421 | competitor for GAM42a | GCC TTC CCA CTT CGT TT | 35 | (Manz et al., 1992) |
| POL740 | *Polaribacter* | CCC TCA GCG TCA GTA CAT ACG T | 35 | (Malmstrom et al., 2007) |
| CF968 | *Bacteroidetes* | GGT AAG GTT CCT CGC GTA | 55 | (Acinas et al., 2015) |
| Opi346 | *Opitutales* | TTC GAA ACT GCT GCC ACC C | 20 | (Reintjes 2017) |
| Cren554 | *Thaumarchaeota* | TTA GGC CCA ATA ATC MTC CT | 20 | (Massana et al., 1997) |
| SAR406-97 | SAR406 clade (*Marinimicrobia*) | CAC CCG TTC GCC AGT TTA | 40 | (Fuchs et al., 2005) |
| DELTA495a | *Deltaproteobacteria* | AGT TAG CCG GTG CTT CCT | 35 | (Loy et al., 2002) |
| cDELTA495a | competitor for DELTA495a | AGT TAG CCG GTG CTT CTT | 35 | (Lücker et al., 2007) |
| DELTA495b | *Deltaproteobacteria* | AGT TAG CCG GCG CTT CCT | 35 | (Loy et al., 2002) |
| cDELTA495b | competitor for DELTA495b | AGT TAG CCG GCG CTT CKT | 35 | (Lücker et al., 2007) |
| DELTA495c | *Deltaproteobacteria* | AAT TAG CCG GTG CTT CCT | 35 | (Loy et al., 2002) |
| cDELTA495c | competitor for DELTA495c | AAT TAG CCG GTG CTT CTT | 35 | (Lücker et al., 2007) |
| SAR324-R-625 | SAR324 clade (“Marine group B”) | CGA AAG ACC CTC CGG | 15 | (Wright et al., 1997) |
| ROS536 | *Rhodobacteraceae* | CAA CGC TAA CCC CCT CCG | 35 | (Brinkmeyer et al., 2000) |

| | | | | |
|---|---|---|---|---|
| SAR11-152R | SAR11 clade | TTAGCACAAGTTTCCYCGTGT | 25 | (Morris et al. 2002) |
| SAR11-441R | SAR11 clade | TACAGTCATTTTCTTCCCCGAC | 25 | (Morris et al. 2002) |
| SAR11-441Rmod | SAR11 clade | TACCGTCATTTTCTTCCCCGAC | 25 | (Gomez-Pereira et al., 2013) |
| SAR11-542R | SAR11 clade | TCCGAACTACGCTAGGTC | 25 | (Morris et al. 2002) |
| SAR11-732R | SAR11 clade | GTCAGTAATGATCCAGAAAGYTG | 25 | (Morris et al. 2002) |
| SAR11-487Rmodif | SAR11 clade | CGGACCTTCTTATTCGGG | 25 | (Gomez-Pereira et al., 2013) |
| SAR11-487_h3 | SAR11 clade | CGGCTGCTGGCACGAAGTTAGC | 25 | (Gomez-Pereira et al., 2013) |

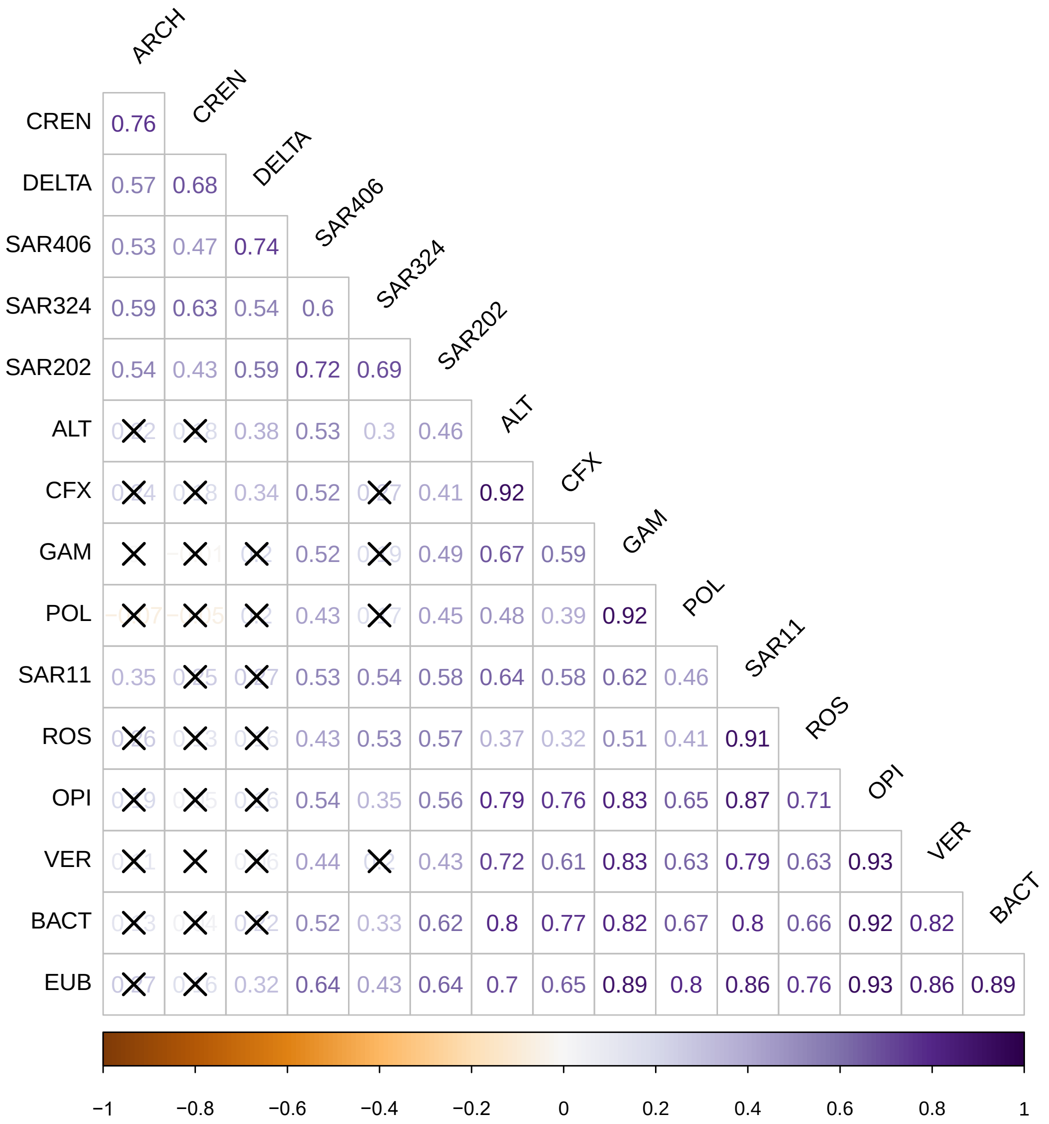


**Figure S1.** Pearsons's correlation coefficient (*r*) plot between cell abundances of the investigated taxonomic groups in the Fram Strait water column. Insignificant correlations are crossed with (X). The number of data pairs was 10. *Bacteria* (EUB), *Archaea* (ARCH), *Alteromonadaceae/Colwelliaceae/Pseudoalteromonadaceae* (ALT). *Bacteroidetes* (BACT). *Chloroflexi* (CFX). *Thaumarchaeota* (THA). *Deltaproteobacteria* (DELTA). *Gammaproteobacteria* (GAM). *Opitutales* (OPI). *Polaribacter* (POL). *Rhodobacteraceae* (ROS). SAR11. SAR202. SAR324. SAR406. *Verrucomicrobiales* (VER).
